# Intermittent hypoxia promotes functional neuroprotection from retinal ischemia in untreated first-generation offspring

**DOI:** 10.1101/2020.02.26.966457

**Authors:** Jarrod C. Harman, Jessie J. Guidry, Jeffrey M. Gidday

## Abstract

Environmental stimuli can promote short- or long-lasting changes in phenotype through epigenetics. Under certain circumstances, induced phenotypes can be passed through the germline to subsequent generations, providing a novel mechanistic basis for disease heritability. In the present study, we tested the hypothesis that repetitively exposing parents to a nonharmful epigenetic stimulus can promote disease resilience in offspring. Male and female mice were mated following brief exposures to mild systemic hypoxia every other day for 16 weeks. Electroretinographic determinations of postischemic function in response to transient unilateral retinal ischemia in their 5-month-old F1 progeny revealed significant resilience to injury relative to animals derived from normoxic control parents. Mass spectrometry identified hundreds of differentially expressed proteins between protected and injured retinae; bioinformatic analyses of the pathways and networks these proteins comprise provided specific mechanistic insights into the molecular manifestation of this injury-resilient phenotype. Thus, epigenetics can modify heritability to promote disease resilience.

## INTRODUCTION

Epigenetics drive cellular responses to stress and environmental change in the form of changes in gene expression (*1*), which in some instances exert long-lasting effects. As examples, early life stress can impact a variety of neuropsychiatric outcomes later in life (*2, 3*). Similarly, cardiovascular disease, cancer, and even aging itself are epigenetically influenced by a myriad of adverse stress stimuli experienced during childhood (*4, 5*). In fact, epigenetic modifications secondary to stress and environmental change can not only affect phenotypes across the lifespan, but can also affect phenotypes in progeny. Accumulating evidence documents the intergenerational and transgenerational passage of epigenetically-induced phenotypes in a number of species (*6–8*), including mammals (*9–11*). That this also occurs in humans, beyond imprinting, is strongly suggested by a broad set of elegantly curated epidemiological data (*12, 13*). The vast majority of research in these aforementioned fields of study has focused on identifying pathological phenotypes and disease susceptibility resulting from previously presented adverse stressors. Rare is the finding that adaptive phenotypes can be induced by nonharmful stressors, despite the likelihood that such beneficial responses actually represent the largely unrecognized other half of the epigenetic lifespan/inheritance fields.

Several decades of preclinical studies in cardiac (*14*) and cerebral (*15*) conditioning medicine consistently confirm that epigenetically-mediated changes in gene expression induced in response to a nonharmful conditioning stimulus promote a transient resilience to acute tissue ischemia. From these and related studies, it is becoming increasingly clear that the magnitude, duration, and frequency of the conditioning stimulus not only dictate whether the induced change in phenotype is adaptive or maladaptive, but also that the frequency, or intermittent nature, of the stimulus proportionally affects the duration of the response (*16–20*). Such findings raise the possibility of leveraging conditioning stimuli to induce, throughout the lifespan, not just deleterious phenotypes, but beneficial epigenetic modifications that promote health and disease resiliency. In turn, repetitive/intermittent conditioning may even induce adaptive epigenetic changes in the germline that promote injury resilience in offspring.

To test the hypothesis that intergenerational epigenetic inheritance occurs in the central nervous system of mammals, we used a well-established mouse model of transient, unilateral retinal ischemia. The world-wide incidence of ischemic retinopathies, including nonproliferative diabetic retinopathy, glaucoma, age-related macular degeneration, and acute ischemia secondary to branch and central retinal artery occlusion, is significant (*21–24*); however, efficacious therapies designed to protect the retina against the consequent blindness and related visual morbidities have yet to be developed. To ensure rigor, we quantified postischemic retinal function by electroretinography in large cohorts. Repetitive systemic hypoxia, which has a human correlate in remote limb conditioning (*25*), served as the epigenetic conditioning stimulus. We also interrogated the intergenerational neuroprotective phenotype by quantitative mass spectrometry of retinae from F1 mice derived from conditioned and nonconditioned parents. Subsequent bioinformatic analyses of these novel protein expression profiles identified diverse signaling pathways/networks that likely underlie this inherited resilience to ischemic injury.

## RESULTS

### Parental Hypoxia Provides Functional Protection Against Retinal Ischemia in Adult Offspring

To assess whether epigenetics-mediated resilience to retinal ischemic injury induced by RHC in the parental generation (*16*) is a phenotype that can be inherited by their first-generation offspring, we used scotopic ERG to measure postischemic retinal function in adult, first-generation (F1) progeny born to RHC-treated parents, and in matched F1 progeny born to matched (untreated) control parents. Retinal ischemia in F1 control mice resulted in a 39% and 27% reduction in a-wave (Fig. 1A) and b-wave (Fig. 1B) amplitudes, respectively, at the highest flash intensity relative to their pre-ischemic baseline responses. In contrast, the extent of a-wave and b-wave amplitude loss was reduced by only 20% and 17%, respectively in the ischemic retinae of mice born to RHC-treated parents. As shown in Figure 1C, when ERG waveform amplitudes of the ischemic retinae were normalized within each animal to their respective contralateral nonischemic retinae for both groups, retinae of mice from F1 controls (n=23) exhibited a 30±4 and 22±4% loss in a-wave and b-wave amplitudes as a result of ischemia, whereas mice from RHC-treated parents only showed an 11±2 (p<0.0001) and 4±3% (p<0.0005) loss, respectively, in a-wave and b-wave amplitudes. That this protective effect resulted from true ischemic resiliency and not any treatment-induced shift in baseline reactivity is borne out by our documentation in the very same animals that no differences in baseline, pre-ischemic a-wave and b-wave amplitudes existed in response to any flash intensity (Supplemental Fig. 1).

**Figure 1.**
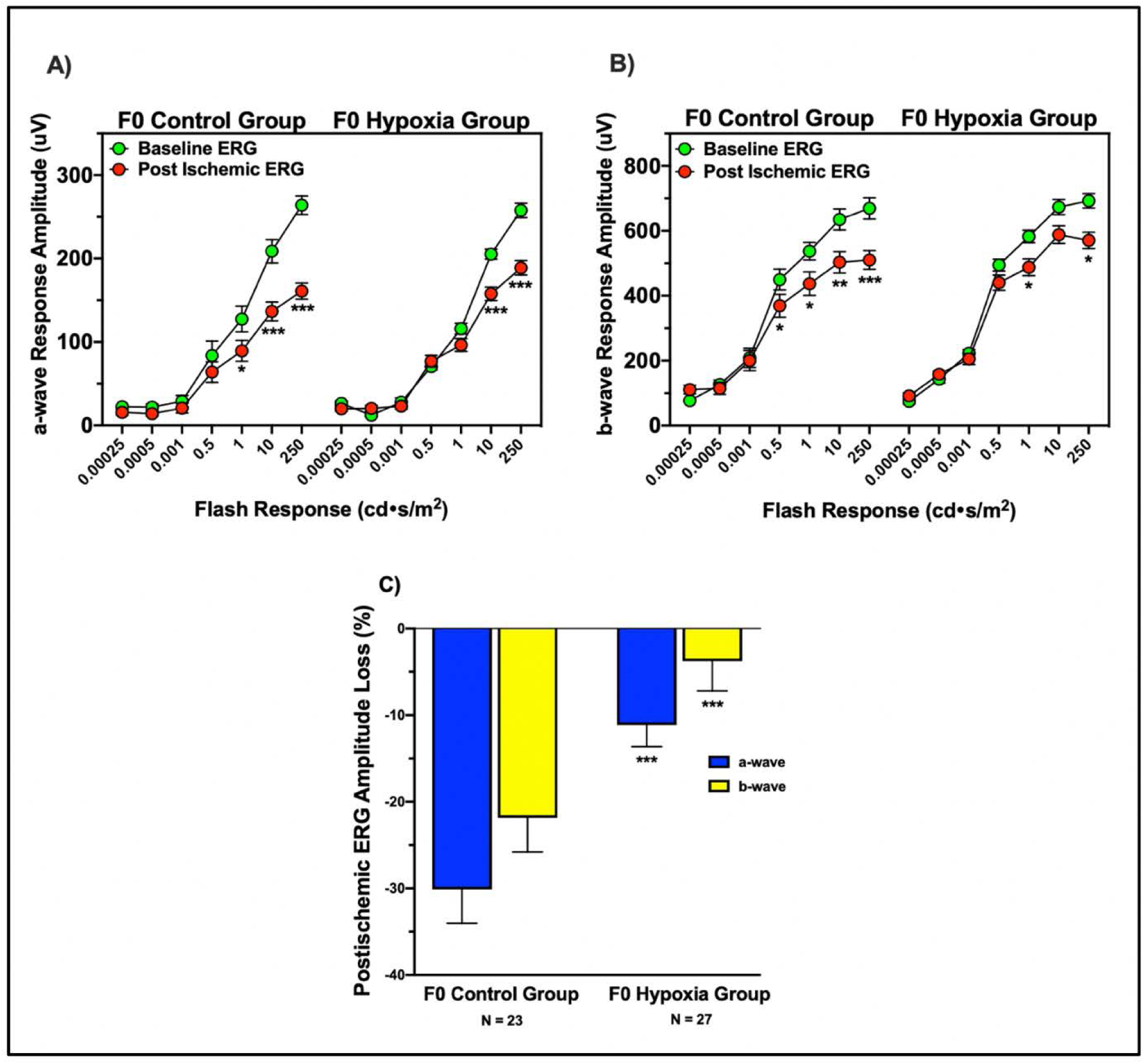
Inherited neuroprotection demonstrated functionally by scotopic electroretinography (ERG). ERG a-wave **(A)** and b-wave **(B)** amplitudes (in microvolts) as a function of increasing light intensity, at pre-ischemic baseline (green) and at 7 days post-ischemia (red) for naïve F1 mice derived from untreated F0 parents (F0 Control Group) and naïve F1 mice derived from F0 parents treated with repetitive hypoxic conditioning prior to mating (F0 Hypoxia Group). **(C)** Normalized reductions in a-wave and b-wave amplitudes in the ischemic eye relative to the nonischemic eye at the highest light intensity (250cd•s/m^2^) in the same F1 Control (n=23) and F1 Hypoxia (n=27) groups. Mean±SEM. *p<0.05-0.005, **p<0.004-0.0005, and ***p<0.0004-0.00005 versus respective Control ERG waveform amplitude at the same light intensity.

### Mass Spectrometry Experimental Design Platform

To probe the mechanistic basis of this intergenerational adaptive phenotype that we documented at the functional level by electroretinography, we performed quantitative, discovery-based proteomic analyses of retinae derived from F1 animals born to RHC-treated parents (F1-*RHC mice) and to normoxic control parents (F1-control mice), selecting animals that were most ‘representative’ of the mean loss of ERG waveform magnitude. Specifically, ischemic and nonischemic retinae from 4 age-matched (22±2 weeks old) males (2 animals derived from control parents and 2 animals derived from RHC-treated parents) were collected 10 days after ischemia (3 days after postischemic ERG). This established the following experimental groups, for which we identified and analyzed their respective differential proteomes:

<u>Comparison 1 (C1)</u>: Ischemic retinae of F1 mice derived from untreated control F0 parents versus nonischemic retinae of F1 mice derived from untreated control F0 parents
<u>Comparison 2 (C2)</u>: Ischemic retinae of F1 mice derived from hypoxic F0 parents versus nonischemic retinae of F1 mice derived from untreated control F0 parents
<u>Comparison 3 (C3)</u>: Ischemic retinae of F1 mice derived from hypoxic F0 parents versus ischemic retinae of F1 mice derived from untreated control parents

These comparisons are illustrated schematically in Figure 2.

**Figure 2.**
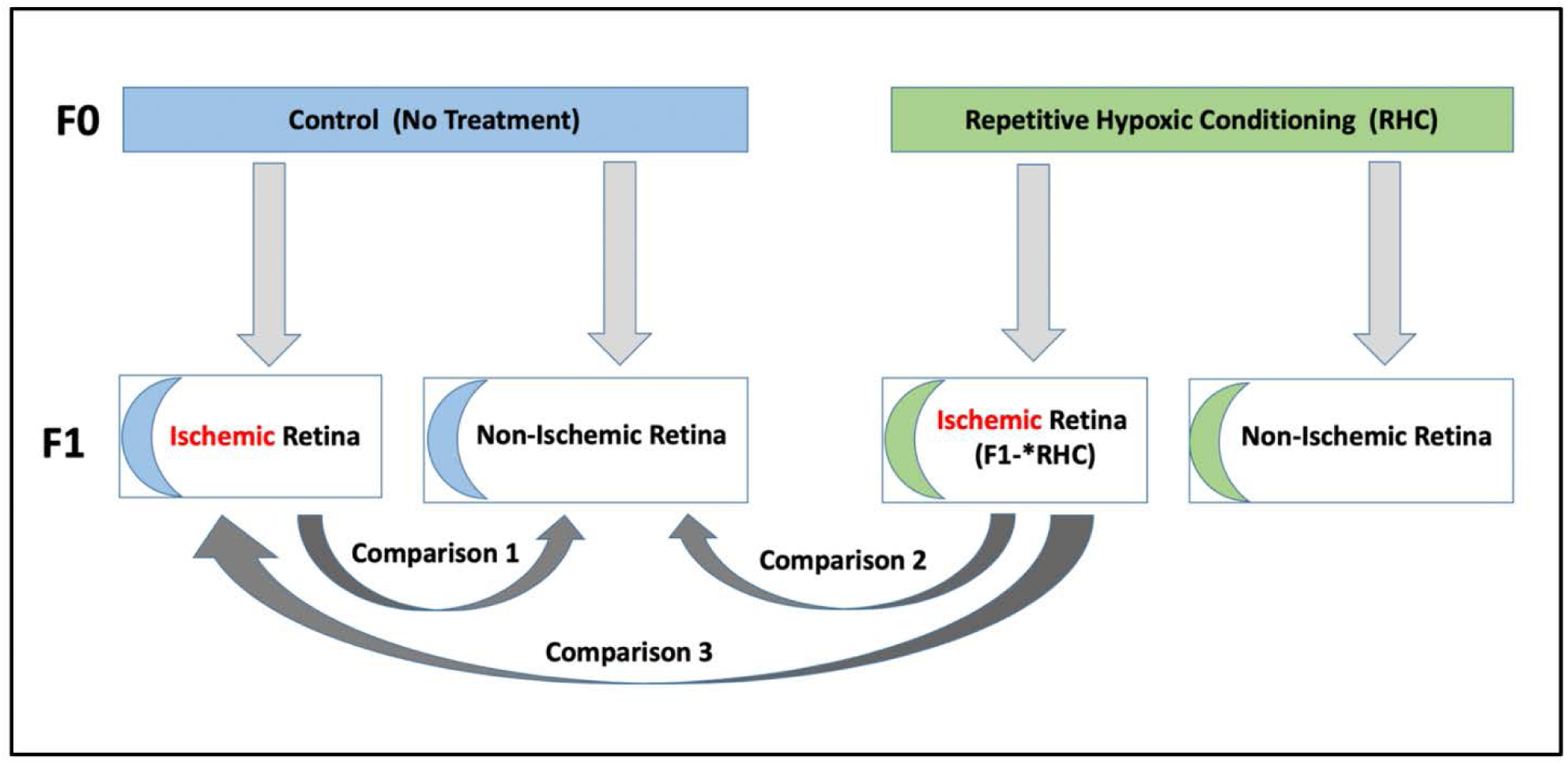
Three Differential Retinal Proteome Comparisons. Shown schematically are the differential proteomes resulting from F0 RHC and F1 ischemia as variables, expressed as relative differences (ratios) or “comparisons” in the manuscript. As such, they serve as a reference platform for describing experimental outcomes.

Overall, 4,175 retinal proteins were identified in our dataset, 4041 with quantitation, which were subsequently loaded into IPA for bioinformatic analyses of a net 1432 ‘analysis-ready’ proteins, after applying “mild” filtering as described above. Table 1 provides the top 10 up-regulated and top 10 down-regulated proteins, based on abundance/fold-change, for each of the three experimental comparisons of interest (see Figure 2) that we will use herein to reference all data displays. In Supplemental Table 1, an expanded list that includes the top 100 differentially expressed proteins for each comparison is provided. Volcano plots generated for each comparison (Suppl. Fig. 2) provide a global quantitative picture of protein distributions, based on both fold change and statistical significance, for each of the 4042 proteins identified. Companion heat maps (Suppl. Fig. 2) highlight the distinct expression profiles for the top 500 differentially expressed proteins in each comparison.

**Table 1.** Top 20 Differentially Expressed Proteins (abundance ratios)

| Comparison 1 |  |  | Comparison 2 |  |  | Comparison 3 |  |  |
| --- | --- | --- | --- | --- | --- | --- | --- | --- |
| GENE ID | PROTEIN NAME | FC RATIO | GENE ID | PROTEIN NAME | FC RATIO | GENE ID | PROTEIN NAME | FC RATIO |
| Serpina3n | Serine protease inhibitor A3N | 10.50 | Ankrd54 | Ankyrin repeat domain-containing protein 54 | 5.10 | Hmgcs2 | Hydroxymethylglutaryl-CoA synthase, mitochondrial | 4.06 |
| Fbln1 | Fibulin-1 | 9.29 | Comtd1 | Catechol O-methyltransferase domain-containing protein 1 | 4.56 | Krt14 | Keratin, type I cytoskeletal 14 | 3.80 |
| Serpina1d | Alpha-1-antitrypsin 1-4 | 6.89 | Hist1h3i;<br>Hist1h3a;<br>Hist1h3g;<br>Hist1h3h | Histone H3.1 | 4.03 | Ankrd54 | Ankyrin repeat domain-containing protein 54 | 3.63 |
| Arg1 | Arginase-1 | 6.16 | Glycam1 | Glycosylation-dependent cell adhesion molecule 1 | 3.97 | Crygs | Beta-crystallin S | 3.51 |
| Cfb | Complement factor B | 6.07 | 4933407H<br>18Rik;<br>Uvssa | UV-stimulated scaffold protein A | 3.87 | Arhgef10 | rho guanine nucleotide exchange factor 10 | 3.40 |
| Serpina3k | Serine protease inhibitor A3K | 6.06 | Dctn6 | Dynactin subunit 6 | 3.71 | Crybb1 | Beta-crystallin B1 | 3.12 |
| Igkc | Ig kappa chain C region | 6.05 | Slc24a4 | Sodium/potassium/calcium exchanger 4 | 3.63 | Grfin | Grfin | 2.90 |
| C1qa | Complement C1q subcomponent subunit A | 5.82 | Hmgcs2 | Hydroxymethylglutaryl-CoA synthase, mitochondrial | 3.45 | Crybb2 | Beta-crystallin B2 | 2.80 |
| Serpina1b | Alpha-1-antitrypsin 1-2 | 5.69 | Cabp5 | Calcium-binding protein 5 | 3.43 | Comtd1 | Catechol O-methyltransferase domain-containing protein 1 | 2.73 |
| Ces1c | Carboxylesterase 1C | 5.55 | Metap1 | Methionine aminopeptidase 1 | 3.28 | Slc2a13 | proton myo-inositol cotransporter | 2.68 |
| Cryge | Gamma-crystallin E | -11.24 | Cryge | Gamma-crystallin E | -38.46 | Serpina1b | Alpha-1-antitrypsin 1-2 | -7.87 |
| C030006K<br>11Rik | UPF0598 protein C8orf82 homolog | -6.41 | Arl6ip1 | ADP-ribosylation factor-like protein 6-interacting protein 1 | -9.09 | Serpina1a;<br>Serpina1d | Alpha-1-antitrypsin 1-1 | -7.64 |
| Ppip5k1 | Inositol hexakisphosphate/diphosphoinositolpentakisphosphate kinase 1 | -6.06 | Cryga | Gamma-crystallin A | -7.82 | Mup11;<br>Mup18;<br>Mup2;<br>Mup19 | major urinary protein 18 | -7.30 |
| Palm2 | paralemmin-2 | -4.20 | C030006K<br>11Rik | UPF0598 protein C8orf82 homolog | -6.21 | Serpina3k | Serine protease inhibitor A3K | -7.14 |
| Crygd | Gamma-crystallin D | -4.18 | Ppip5k1 | Inositol hexakisphosphate and diphosphoinositolpentakisphosphate kinase 1 | -5.81 | Tgfb1 | Transforming growth factor-beta-induced protein ig-h3 | -6.99 |
| Thoc5 | THO complex subunit 5 homolog | -4.12 | Thoc5 | THO complex subunit 5 homolog | -5.62 | Arl6ip1 | ADP-ribosylation factor-like protein 6-interacting protein 1 | -6.58 |
| Cwc15 | spliceosome-associated protein CWC15 homolog | -3.78 | Tubb6 | Tubulin beta-6 chain | -5.29 | Ces1c | Carboxylesterase 1C | -6.58 |
| Fabp5 | Fatty acid-binding protein, epidermal | -3.48 | 1110037F<br>02Rik;<br>Virma | protein virilizer homolog | -4.57 | Serpina3n | Serine protease inhibitor A3N | -6.41 |
| Grfin | Grfin | -3.46 | Cwc15 | spliceosome-associated protein CWC15 homolog | -3.85 | Alb | Serum albumin | -6.02 |
| Bfsp2 | Phakinin | -3.42 | Slc25a46 | Solute carrier family 25 member 46 | -3.81 | Apoa4 | Apolipoprotein A-IV | -5.85 |

### The Postischemic Response

Comparing the proteome of ischemic retinae in control F1 mice derived from untreated parents to nonischemic retinae in control F1 mice derived from untreated parents – what we term herein as the *‘Control Ischemic Response’*, or Comparison 1 (C1) (Table 1 and Suppl. Table 1) – identified the differentially expressed proteins that defined the phenotype of this tissue 10 days post-ischemia. This included 411 proteins expressed in higher abundance, and 80 proteins expressed in lower abundance (491 differentially expressed proteins in total). 233 of the 411 differentially upregulated proteins, and 38 of the 80 differentially downregulated proteins were not differentially expressed in response to ischemia in mice derived from RHC-treated parents. Bioinformatic analyses of the dataset characterizing the *control ischemic response* yielded the following (IPA-defined) top 5 enriched “Canonical Pathways” (including the number of differentially expressed proteins out of the total number of proteins in the pathway, along with the p-value of overlap) that are enriched in proteins that were differentially expressed in response to ischemia: ‘Acute Phase Response Signaling’ (26/179 proteins, p=5.5E-21), ‘LXR/RXR Activation’ (17/121 proteins, p=7.0E-14), ‘FXR/RXR Activation’ (17/126 proteins, p=1.4E-13), ‘Coagulation System’ (9/35 proteins, p=2.0E-10), and ‘RhoGDI Signaling’ (14/180 proteins, p=3.1E-8). The top five ‘downstream’ “Molecular and Cellular Functions” (including the number of proteins in the pathway, and the p-value range) defining the *control ischemic response* were: ‘Cellular Movement’ (117 proteins, p-value=5.3E-6→4.0E-23), ‘Cellular Assembly and Organization’ (111 proteins, p=4.2E-6→1.7E-18), ‘Protein Synthesis’ (68 proteins, p=4.1E-6→2.4E-17), ‘Cell-to-Cell Signaling and Interaction’ (97 proteins, p=5.0E-6→1.5E-16), and ‘Cellular Compromise’ (47 proteins, p=3.6E-06→1.6E-16). The top 5 “Upstream Regulators” (with p-values) of the *control ischemic response* – the upstream molecules that are responsible for the regulation patterns we see in this particular response – were: the multi-functional regulator transforming growth factor-β1 (TGF-β1; p=2.3E-18), prolactin (PRL; p=1.5E-16), the tumor suppressor p53 (TP53; p=4.8E-15), the inflammatory cytokine interleukin-6 (IL6; p=1.7E-12), and the chromatin-regulating high mobility group nucleosomal binding domain containing protein 3 (Hmgn3; p=2.2E-12).

### Parental Hypoxia Modifies the Postischemic Response

The retinal proteomic response to ischemia in F1 mice born to RHC-treated F0 parents, relative to nonischemic retinae in control F1 mice derived from untreated parents – what we term the *‘RHC Ischemic Response’* or Comparison 2 (C2) – differed in many unique and significant ways from the aforementioned *control ischemic response* of F1 mice born to untreated F0 parents (C1). In total, 402 proteins were found to be significantly differentially expressed, with 314 exhibiting higher abundances, and 88 exhibiting lower abundances, relative to nonischemic retina. Moreover, 141 of the 314 differentially upregulated proteins, and 42 of the 88 differentially downregulated proteins were not found differentially expressed in response to ischemia in untreated mice (the *control ischemic response*) (Suppl. Table 1). Bioinformatic analyses of this *RHC ischemic response* revealed the top five enriched canonical pathways to be ‘CXCR4 Signaling’ (11/167 proteins, p=2.2E-7), ‘Ephrin B Signaling’ (7/72 proteins, p=2.8E-6), ‘Ephrin Receptor Signaling’ (10/180 proteins, p=3.7E-6), ‘Cardiac Hypertrophy Signaling’ (11/240 proteins, p=7.7E-6), and ‘Histamine Degradation’ (4/17 proteins, p=1.2E-5). The top 5 Molecular and Cellular Functions were ‘Cellular Assembly and Organization’ (48 proteins, p-value=8.6E-3→1.2E-6), ‘Cellular Function and Maintenance’ (47 proteins, p=8.6E-3→1.2E-6), ‘Amino Acid Metabolism’ (14 proteins, p=7.2E-3→3.8E-6), ‘Small Molecule Biochemistry’ (43 proteins, p=8.6E-3→3.8E-6), and ‘Cell-to-Cell Signaling and Interaction’ (35 proteins, p=8.6E-3→6.6E-5). The top 5 Upstream Regulators of the RHC ischemic response were: TP53 (p=1.1E-5), amyloid beta precursor protein (APP; p=3.1E-5), cytochrome P450 oxidoreductase (POR; p=2.1E-4), mitogen-activated protein kinase-1 (MAPK1; p=3.4E-4), and microRNA miR-1228-5p (p=3.8E-4).

It is clear that, at the level of individual proteins and pathways, parental RHC fundamentally changed the response to ischemia in their offspring. None of the top five Canonical Pathways was shared between C1 and C2, and the only top five Upstream Regulator they had in common was TP53. With respect to the top five Molecular and Cellular Functions, only the ‘Cellular Assembly and Organization’ pathway and the ‘Cell-to-Cell Signaling and Interaction’ pathway were shared.

To further probe these datasets, we performed additional bioinformatic analyses of only the proteins that were differentially expressed in C1, but not C2, as well as, conversely, only the proteins that were differentially expressed in C2, but not C1. Omitting from the analysis the proteins that were differentially expressed in one Comparison but not the other provided more mechanistic insights into each response, while providing additional confirmation of the original findings for each Comparison despite the inclusion of redundant proteins. In terms of IPA’s Core Analyses, and the proteins that were only differentially expressed in C1, but not in C2, the top 5 “Canonical Pathways” were: ‘Acute Phase Response Signaling’ (23/179 proteins, p=7.5E-18), ‘LXR/RXR Activation’ (15/121 proteins, p=7.8E-12), ‘FXR/RXR Activation’ (15/126 proteins, p=1.4E-11), ‘Coagulation System’ (9/35 proteins, p=1.4E-10), and ‘Actin Cytoskeleton Signaling’ (13/218 proteins, p=1.3E-6). In turn, the top five ‘downstream’ “Molecular and Cellular Functions” were: ‘Protein Synthesis’ (63 proteins, p=2.5E-5→1.1E-16), ‘Cellular Movement’ (105 proteins, p-value=5.9E-5→2.8E-16), ‘Cellular Compromise’ (41 proteins, p=1.1E-05→1.2E-13), ‘Free Radical Scavenging (35 proteins, p=1.6E-5→7.3E-13), and ‘Molecular Transport’ (86 proteins, p=5.1E-5→2.4E-11). Finally, the top 5 “Upstream Regulators” were: the multi-functional regulator transforming growth factor-β1 (TGF-β1; p=9.9E-15), the chromatin-regulating high mobility group nucleosomal binding domain containing protein 3 (Hmgn3; p=1.8E-12), prolactin (PRL; p=4.6E-12), the tumor suppressor p53 (TP53; p=3.4E-10), and the cell proliferation regulator and Ras-based GTPase KRAS (p=4.0E-10). Of note, 4/5 canonicals, 3/5 molecular and cellular functions, and 4/5 upstream regulators were identical, even after removing the 220 proteins from the C1 dataset that were not differentially expressed in C2.

With respect to the proteins that were only differentially expressed in C2, but not in C1, the top 5 “Canonical Pathways” were ‘Endocannabinoid Developing Neuron Pathway’ (6/115 proteins, p=2.5E-4), ‘CXCR4 Signaling’ (7/167 proteins, p=2.9E-4), ‘Antiproliferative Role of Somatostatin Receptor 2’ (5/77 proteins, p=3.0E-4), ‘Ephrin Receptor Signaling’ (7/180 proteins, p=4.5E-4), and ‘BMP Signaling Pathway’ (5/85 proteins, p=4.7E-4). The top 5 Molecular and Cellular Functions were: ‘Cell-to-Cell Signaling and Interaction’ (33 proteins, p=8.1E-3→5.2E-5), ‘Amino Acid Metabolism’ (9 proteins, p=7.6E-3→5.6E-5), ‘Small Molecule Biochemistry’ (40 proteins, p=8.1E-3→5.7E-5), ‘Cellular Development’ (46 proteins, p=8.6E-3→5.7E-5), and ‘Molecular Transport’ (46 proteins, p=8.1E-3→1.6E-4). And the top 5 Upstream Regulators of the RHC ischemic response were: TP53 (p=6.0E-6), microRNA miR-450b-3p (p=5.4E-5), cytochrome P450 oxidoreductase (POR; p=9.6E-5), the DNA-binding homeobox protein EN1 (p=1.3E-4), and paired box transcription factor family member PAX3 (p=1.72E-4). In this case, removing the 219 proteins from C2 that were not differentially expressed in C1 yielded 3/5 new top-ranked canonicals, 2/5 new molecular and cellular functions, and 3/5 new upstream regulators.

### Two Distinct Postischemic Retinal Proteomes

Directly comparing postischemic retinal proteomes of mice from RHC-treated parents to that of mice from untreated parents – which we term the *‘RHC vs Control Ischemic Response’* or Comparison 3 (C3) – provides another distinct way to probe the mechanistic basis of the ischemia-resilient phenotype. In total, 266 proteins were found to be significantly differentially expressed in this dataset, with 73 proteins exhibiting higher abundances, and 193 proteins exhibiting lower abundances (Table 1 and Suppl. Table 1). These differences support the general conclusion that, at the timepoint examined, parental RHC results in an overall suppression of ischemia-induced changes in protein expression. Many of these downregulated proteins (e.g., integrins, G-protein-coupled receptors, and receptor tyrosine kinases) serve as ‘molecular switches’ that activate or inhibit signaling pathways. The top 5 Canonicals representing this comparison were: ‘Acute Phase Response Signaling’ (22/179 proteins, p=6.9E-22), ‘LXR/RXR Activation’ (16/121 proteins, p=1.1E-16), ‘FXR/RXR Activation’ (16/126 proteins, p=2.0E-16), ‘Coagulation System’ (9/35 proteins, p=1.1E-12), and ‘Complement System’ (6/37 proteins, p=1.4E-7). The top five “Molecular and Cellular Functions” defining C3 were: ‘Cellular Compromise’ (39 proteins, p=1.3E-4→1.4E-17), ‘Cellular Movement’ (69 proteins, p=1.3E-4→2.6E-15), ‘Protein Synthesis’ (44 proteins, p=1.3E-4→9.8E-15), ‘Lipid Metabolism’ (56 proteins, p=1.3E-4→1.6E-11), and ‘Molecular Transport’ (60 proteins, p=1.3E-04→1.6E-11). Finally, the top “Upstream Regulators” for this *RHC vs control ischemia* comparison were: Hmgn3 (p=6.3E-15), TP53 (p=2.4E-13), PRL (p=9.5E-13), IL6 (p=3.2E-12), and the photoreceptor-specific transcription factor known as cone-rod homeobox protein (CRX; p=2.6E-10).

We performed a ‘comparison analysis’ of the differentially expressed proteins across all three experimental groups/comparisons, and, based on activation z-scores (which are used to predict activation or inhibition, with scores ≥ ±2 indicative of statistical significance), we ranked the top 30 Canonical pathways, Upstream Regulators, and Molecular and Cellular (Biological) Functions as variables (Figure 3). Examination of the z-score–based ‘heat map’ values for a given pathway, regulator, and function reveal many, sometimes very robust, differences between the three experimental groups/comparisons. For example, an upregulation of canonical integrin signaling defines the response to ischemia in control mice (C1), but less so in *RHC mice (C2), to the extent that, when comparing ischemic retinae from the two groups directly to one another (*RHC vs control ischemia*, C3), the overall reduction in integrin signaling that occurs in the ischemic retina as a result of parental RHC is now evident. The same conclusion applies to the canonical signaling pathways for synaptogenesis, Rho family GTPases, actin cytoskeleton, Rac, LXR/RXR activation, paxillin, leukocyte extravasation, acute phase response, cdc42, Nrf2, and RhoA, to name a few. These comparison analyses also reveal that simply identifying the top canonicals or other pathways/networks for a given data set, as we did above for C1-C3, does not concomitantly provide information about the directional changes of the differentially expressed proteins therein. As a poignant example, ‘Acute Phase Response (APR) Signaling’ was identified as the highest ranked canonical for both C1 and C3, but as revealed by our comparison analysis, overall this pathway was robustly upregulated in control mice, whereas in *RHC mice, overall this pathway was robustly downregulated.

**Figure 3.**
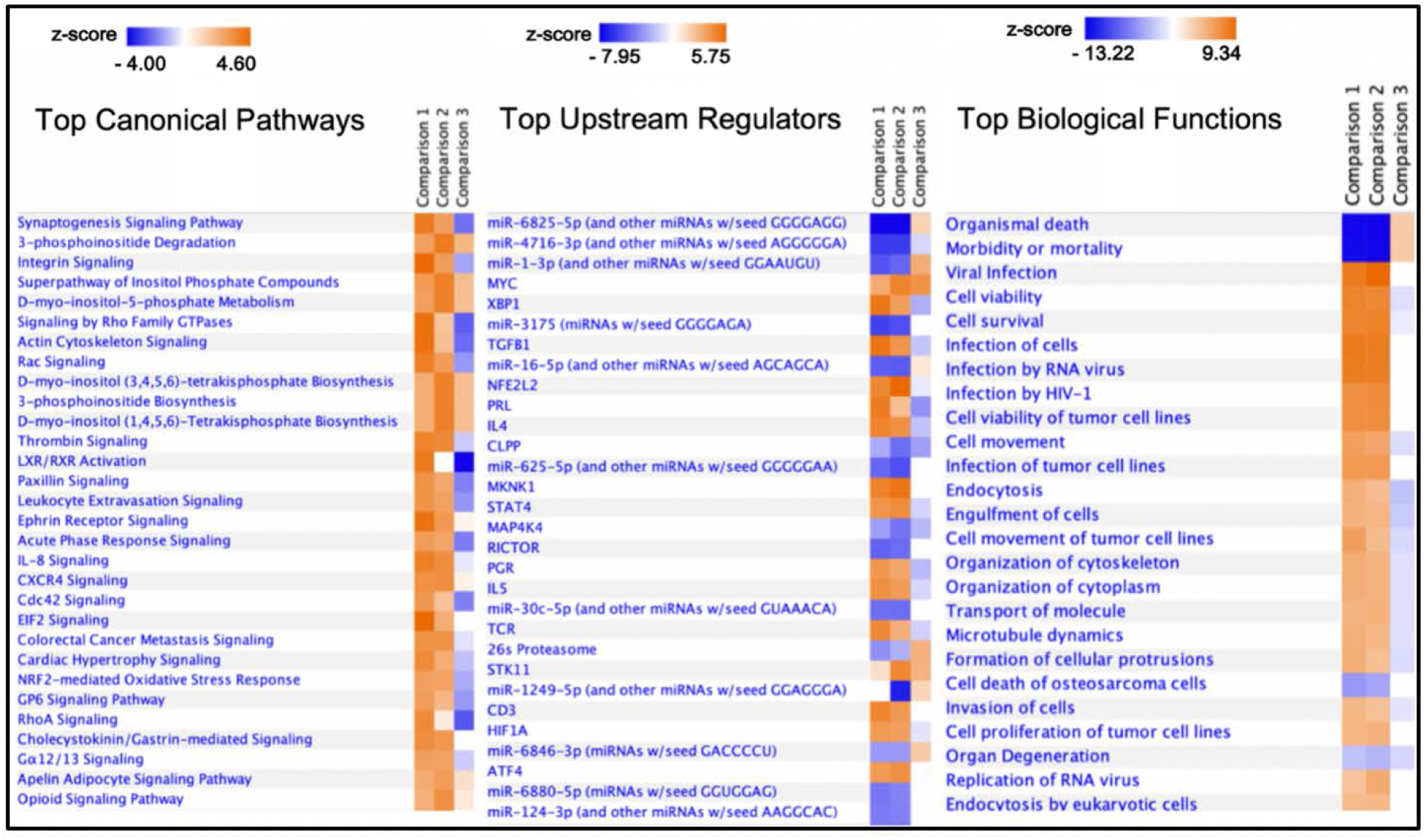
Top canonical pathways, upstream regulators, and biological functional enrichments, by experimental comparison. IPA core analyses were performed for each of the three individual comparisons, and then subsequently compared with each other to yield the most enriched canonical pathways (left), upstream transcriptional regulators (center), and biological/molecular functions (right), with z-score-based heat maps for each pathway, regulator, and function. Only canonicals with a ‘z-score’ of greater than ± 2.0 (=95% confidence interval) are shown. Positive z-scores (orange) indicate activation/upregulation of the pathway, and negative z-scores (blue) indicate suppression/downregulation of the pathway, for each comparison.

In addition to probing the upstream regulators predicted to be driving the observed expression changes, and the downstream biological functions they influence, protein-specific insights into the mechanisms by which parental RHC augments retinal resilience in their offspring can be gleaned from bioinformatically probing the differentially expressed proteins that comprise a given canonical pathway or biological function – for each of the three experimental comparisons – with respect to the magnitude of the respective protein expression changes therein, the location of these differentially expressed proteins in the respective pathway/network, their functional relationship to other differentially expressed proteins, and, as mentioned for the Acute Phase Response Signaling canonical above, the respective direction of their expression changes. We provide three examples below that are illustrative of the mechanistic power of such deeper analyses.

### Actin Cytoskeleton Signaling

Shown schematically in Figure 4 is the actin cytoskeleton signaling network canonical for C3, along with a heat map that displays how each protein in the network is affected under all three experimental conditions (C1-C3). Actin is an abundant, multi-functional protein that forms a complex cytoskeletal network integral to key dynamic processes such as cell motility, intracellular transport, axon guidance, as well as endocytosis and exocytosis; many signal transduction systems use the actin cytoskeleton as a subcellular localization scaffold. Playing central roles in regulating these processes are members of the Rho family of small GTPases, including RhoA and Rac (Ras superfamily members), cdc42 (cell division cycle 42), and ROCK (Rho-associated protein kinase), which, after activation by various classes of transmembrane receptors (i.e., integrin receptors, receptor tyrosine kinases, and G protein-coupled receptors), integrate and transmit signals to downstream effector proteins involved in orchestrating cytoskeletal dynamics. Analyses of this canonical network for C3, and the expression changes that define it, by comparison, in heat map and tabular form (Fig. 4), in conjunction with the corresponding networks for C1 and C2 (Suppl. Figs. 3-upper and 3-lower, respectively), reveal how the signaling responsible for ischemia-triggered polymerization and destabilization of actin, and the assembly of focal adhesions and complexes, is countered by parental RHC. Mechanistically, the expression changes we measured predict that these pathological changes are attenuated, neutralized, or even reversed in the ischemic retinae of *RHC mice as a result of overall increases in the postischemic expression of integrin subunit β1 (by 1.4-fold), RhoA (by 1.9-fold), the adaptor protein BAIAP2 (by 1.2-fold), and the adhesion/contact proteins paxillin [PXN] and talin 1 [TLN] (by 2.3-fold 1.3-fold, respectively. Concomitantly, RHC led to overall decreases in the postischemic expression of several components of the actin-related protein 2/3 complex [ARP2/3] (including the actin-related binding proteins 2 and 3 [ARP2, ARP3] and subunit 1B [ARPC1B]), as well as the actin binding protein filamin-A [FLNA]), the spectrin family protein actinin alpha 4 [ACTN4] (by 1.3-fold), the adherens junction-forming Ras GTPase-activating-like protein IQGAP1 (by 1.4-fold), the adhesion contact protein profilin 2 [PFN] (by 1.3-fold), the F-actin binding protein vinculin [VCL], and the actin filament organizing protein cofilin [CFL1 and CFL2] (by 1.2- and 1.3-fold, respectively). Additional effectors of the RHC-protective phenotype include reductions in the expression of filamentous actin (F-actin) and several myosin light chain [MLC] isoforms, as well as much lower circulating levels of fibronectin [FN1], thrombin [F2], and kinogens like bradykinin that drive cytoskeletal pathobiology. Other top canonicals (Figure 3) related to cytoskeletal dynamics, and which exhibit contrasting activation/inhibition for C1/C3, include ‘RhoA Signaling’, ‘Rac Signaling’, ‘Cdc42 Signaling’, ‘Paxillin Signaling’, and ‘Integrin Signaling’. Similarly, related top Biological Functions (Figure 3) include ‘Cell Movement’, ‘Organization of Cytoskeleton’, ‘Organization of Cytoplasm’, ‘Microtubule Dynamics’, and ‘Formation of Cellular Protrusions’. In addition, directionally similar and statistically significant expression changes were also observed with respect to RHC-associated decreases in ARP2/3, IQGAP1, ERM, MLC, and CFL in two other related canonicals – RhoGTPase Family and RhoGDI Signaling – for C3 (data not shown), all of which underscore the critical role these cytoskeleton proteins likely play in manifesting the ischemia-resilient phenotype.

**Figure 4:**
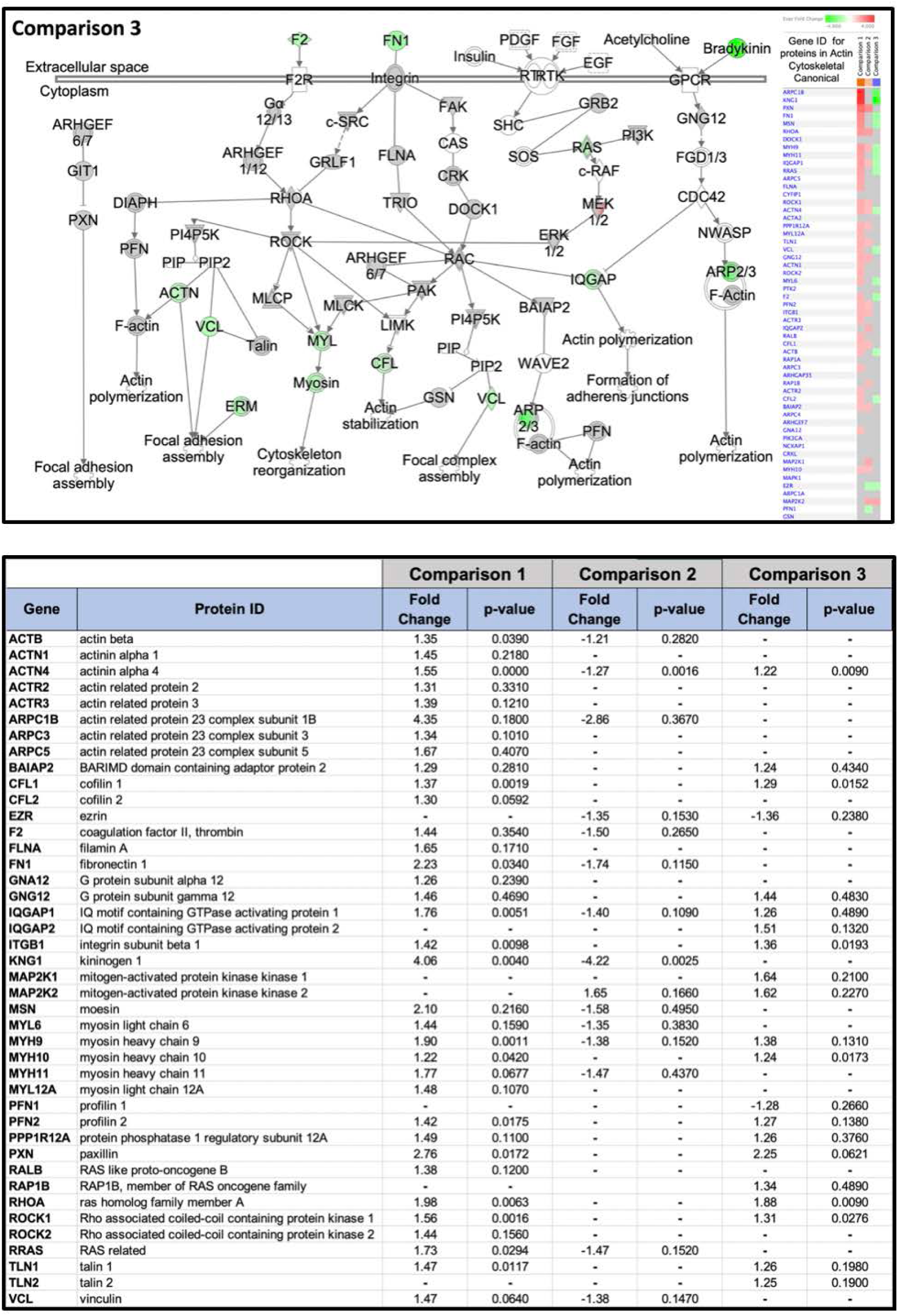
Parental RHC suppresses postischemic ‘Actin Cytoskeleton Signaling’ in F1 retinae. Axon guidance, cell motility, and a myriad of other important cellular functions are directed by actin remodeling. This complex regulatory signaling is directed by many members of the Rho family of small GTPases. Displayed is IPA’s actin cytoskeleton signaling pathway for Comparison 3 (see Supplemental Figure 5 for Comparisons 1 and 2 for this same canonical), with differentially expressed proteins colored red (upregulated) or green (downregulated), based on fold change. The heatmap shows the directional fold-change, by comparison, for each protein in the canonical, ranked by expression fold-change for C1. Note that many of the same proteins that are upregulated in mice from untreated parents (shades of red, Comparison 1, and red-grey in Comparison 2) are downregulated in mice from RHC-treated parents (shades of green, Comparison 3). A tabulated version of this canonical is provided with the fold changes and p-values for each protein in the canonical and each respective comparison.

### Acute Phase Response Signaling

The advent of single-cell proteomics will ultimately provide even more elegant insights into how a given therapeutic affords protection against pathology at the level of specific retinal neurons, Müller cells, and even the retinal and choroidal vasculature. That said, there are some advantages to preclinical sampling of whole, nonperfused tissues; with the resultant ‘capture’ of intravascular proteins, a broader, integrated understanding is gained regarding the many ways injury resilience is ultimately manifested at the tissue level. Indeed, the Coagulant System was a top-ranked canonical for both C1 and C3, and the Complement System was in the top 5 canonicals for C3. A closer examination of the ‘acute phase response’ (APR) to injury, the top canonical for both C1 and C3, serves to emphasize this point. The APR refers to a systemic, innate immune response involving the hepatic release of hundreds of positive and negative acute phase proteins designed to reestablish homeostasis and promote healing following injury (*26*). Figure 5 schematically shows the differentially expressed proteins participating in this inflammatory response for the *RHC-to-control ischemic response* comparison (C3), again alongside a companion heat map showing how each protein in the pathway is affected across all three experimental conditions/comparisons. Also provided in tabular form are the fold changes and respective p-values for the expression differences of all of the proteins in the canonical, sorted by comparison.

**Figure 5.**
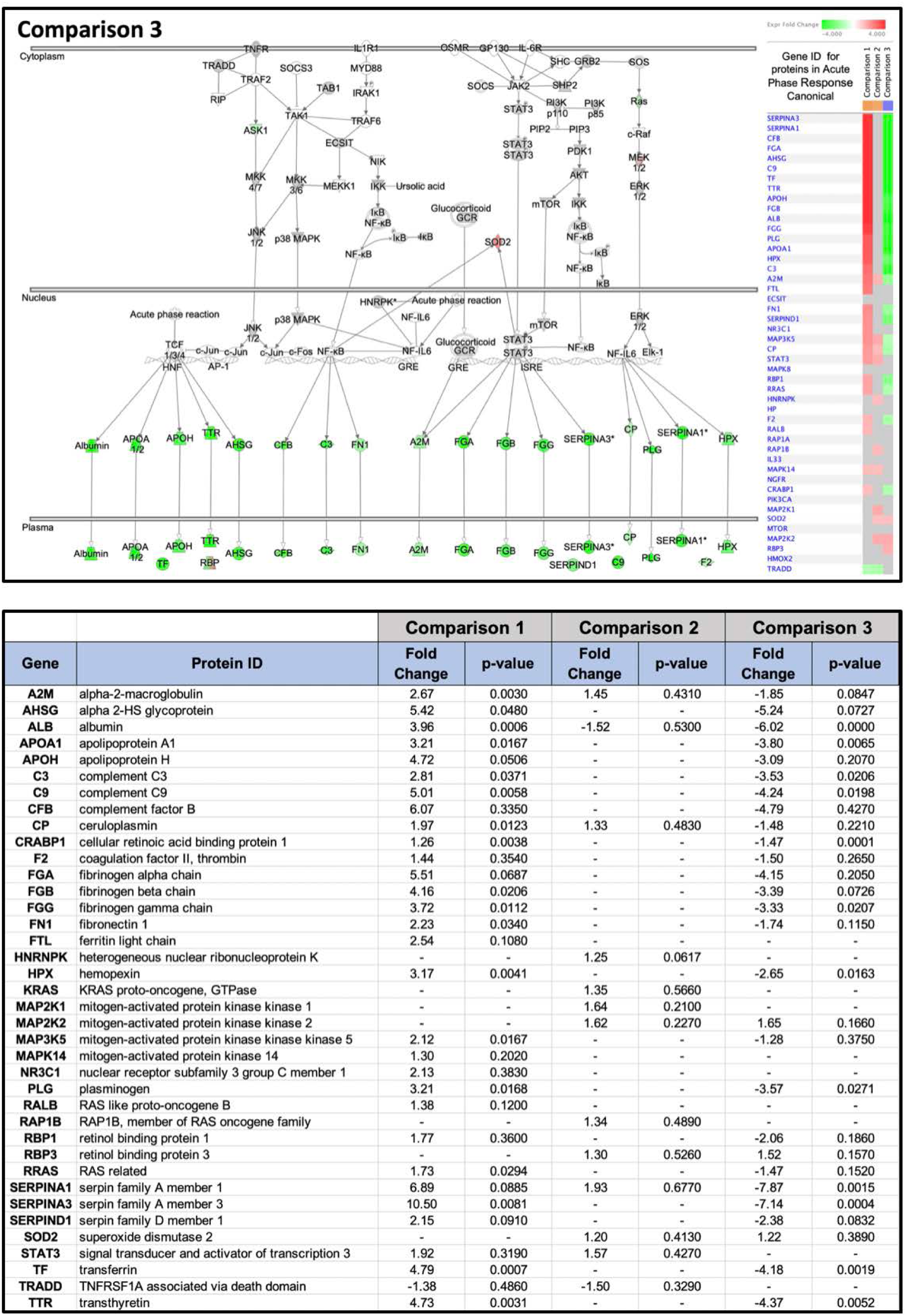
Parental RHC suppresses postischemic ‘Acute Phase Response’ in F1 Retinae. The acute phase response pathway represents systemic pro-inflammatory responses triggered by tissue injury that modify immune function, metabolism, oxidative stress, etc. Displayed is IPA’s version of this pathway for Comparison 3 (see Supplemental Figure 6 for Comparisons 1 and 2 for this same canonical), with differentially expressed proteins colored red (upregulated) or green (downregulated), based on fold change. The heatmap shows the directional fold-change, by comparison, for each protein in the canonical, ranked by expression fold-change for C1. Note that many of the same proteins that are upregulated in mice from untreated parents (shades of red, Comparison 1 are downregulated in mice from RHC-treated parents (shades of green, Comparison 3), reflecting the robust inhibition of this component of ischemic injury as a result of F0 RHC treatment. Note the intra- and inter-cellular distribution (nucleus vs cytoplasm vs plasma) of the proteins in this systemic canonical. A tabulated version of this canonical is provided with the fold changes and p-values for each protein in the canonical and each respective comparison.

Note, the robust downregulation of almost all of the major proteins comprising this particular response to ischemia as a result of parental RHC, including a 3.1- to 3.8-fold downregulation of the A1, A2, and H apolipoprotein transporters, a 4.4-fold downregulation in transthyretin [TTR] expression (which among other functions transports retinol [Vitamin A] in the plasma by associating with retinol binding protein [RPB]), a 1.9- to 5.2-fold downregulation in the levels of the plasma protease regulators alpha-2-HS-glycoprotein [AHSG], alpha-2-macroglobulin, and hemopexin [HPX], a 3.5- to 4.8-fold downregulation in selected complement factors [CFB, C3, C9], a 1.7- to 4.2-fold downregulation in selected coagulation and extracellular matrix regulators (fibronectin [FN1], the alpha [FGA], beta [FGB], and gamma [FGG] subunits of fibrinogen [FGB], plasminogen [PLG], and prothrombin [F2]). Of note, FN1, which serves as a ligand for integrin membrane receptors that modulate actin cytoskeleton signaling and remodeling, as well as plays roles in protein chaperoning, binding, and cell activation in the plasma, differed in expression nearly 4-fold between C1 and C3. Additionally, we found a 2.4- to 7.9-fold downregulation in the selected serine protease inhibitors α1-anti-trypsin [SERPINA1], α1-antichymotrypsin [SERPINA3], and heparin cofactor 2 [SERPIND1]. Conversely, the expression of MnSOD (SOD2), the mitochondrial enzyme that clears superoxide, and thus plays a central role combating ischemia-related oxidative stress and a proinflammatory cytokine burden, is upregulated in the *RHC ischemic retina. These changes contrast those defining the *control ischemic response* for this pathway (C1, Suppl. Figure 4), which is characterized by the robust, 2- to 10-fold, significant upregulation of all of these same pathway proteins (except SERPINA1), as well as others in this canonical, secondary to increases in upstream glucocorticoid receptor activation, p38MAPK and Rac (Suppl. Fig. 4-upper). In turn, the effect of parental RHC (C2) is so robust with regard to inhibiting this overall response systemically that, bioinformatically speaking, the acute phase response does not register as activated (Suppl. Fig. 4-lower), and none of the APR pathway proteins exhibited significant changes in these retinae – at 10 days postischemia. The following are two of many examples that underscore how robustly parental RHC dampened the APR when using C3 as a reference. The proinflammatory cytokine TGF-β1, which was identified as one of the top-ranking upstream regulators in C1 with a z-score of 5.3, was expressed at nearly 7-fold lower levels in C3 (z-score of −1.8), a difference which was also confirmed by our MS results with respect to protein abundance. Additional immune-related differences in protein expression between C1 and C3 are evident in the heatmap differences for a number of biological functions listed in Figure 3, including cellular mobilization, cell invasion, and endocytosis/engulfment of cells.

### Visual Phototransduction

The process of converting photons of light into an electrical signal in the retina, known as visual phototransduction, is what we recorded noninvasively in control and *RHC mice by scotopic (dark-adapted) flash electroretinography; we subsequently quantified at distinct flash intensities the a-wave and b-wave components of the resultant mass potential, which originate from the activity of photoreceptors and cells of the inner retina, respectively, as functional metrics of retinal ischemic injury and resilience (Figure 1). Our MS analysis of ischemic retinae from F1 control mice (C1) and F1-*RHC mice (C2) revealed many rod-specific proteins that, relative to nonischemic retinae, were differentially expressed in each cohort (Figure 6), thereby providing a molecular basis for the epigenetically-mediated functional protection we documented. As evident from the figure, companion heat map, and table of expression differences for the proteins in the canonical, the *control ischemic response* (C1) (Fig. 6, upper panel) is defined by a significant, 2.5-fold reduction in rhodopsin [OPSIN] expression (as well as a 1.5-fold reduction in short wave sensitive opsin-1 [OPN1SW]), a 1.5-fold reduction in regulator of G-protein signaling 9-binding protein [RGS9BP/R9AP] expression, and a 1.2-fold reduction in transducin-α [GNAT1] expression (the subunit that contains the GTP binding site), relative to nonischemic retinae. Also defining this response are 1.6- and 1.8-fold increases in the expression of transducin-β subunits GNB1 and GNB3 (the subunits regulating GTPase activity), and a 1.3-fold increase in S-arrestin [SAG].

**Figure 6.**
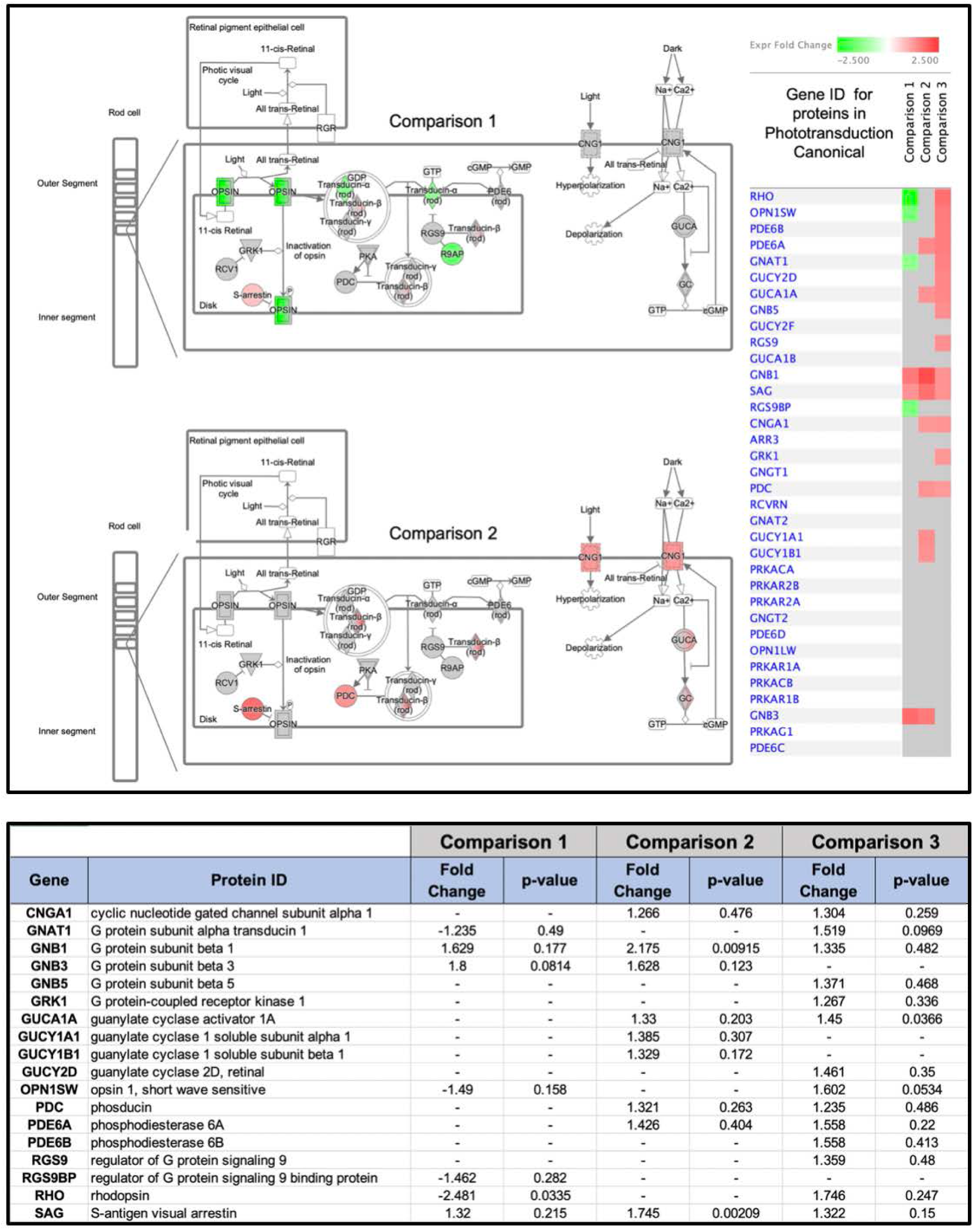
Parental RHC reverses ischemia-induced changes in rod phototransduction pathway proteins. Displayed is IPA’s version of the phototransduction canonical pathway for Comparison 1 (upper panel) and Comparison 2 (lower panel) (see Supplemental Figure 7) for Comparison 3 for this same canonical). Proteins and their isoforms responsible for generating an electrical response to light are arranged anatomically relative to the rod outer segment (the section of the photoreceptor containing disks full of the photosensitive pigment rhodopsin and other molecules involved in the visual transduction process), as well as the channels and molecular mediators of the so-called ‘dark current’ that maintain the rod in a depolarized state in the absence of light. The heatmap shows the directional fold-change, by comparison, for each protein in the canonical, ranked by expression fold-change for C1. Note that many of the proteins that are downregulated (green) in the ischemic retinae of F1 mice from control parents (Comparison 1) are either no longer downregulated or even upregulated (red) in the ischemic retinae of F1 mice from RHC-treated parents (Comparison 2). A tabulated version of this canonical is provided with the fold changes and p-values for each protein in the canonical and each respective comparison.

In contrast, the *RHC ischemic response* (C2) (Fig. 6, lower panel) reveals in striking fashion how the rod proteome is completely modified by parental RHC. No longer are the aforementioned downregulated proteins affected. Rather, the expression of one of these is now upregulated (transducin-α [GNAT1], by 1.2 fold), and the expression of several other photoreceptor proteins that were unaffected by ischemia are now significantly upregulated: specifically, phosphodiesterase-6 α [PDE6A] by 1.4 fold, and the phosphoprotein phosducin [PDC], by 1.3 fold. Of note, the ischemia-induced increases in transducin-β and S-arrestin expression were significantly increased in *RHP mice, even more so than in untreated mice, consistent with the notion that augmenting their expression may be an endogenous response on the part of the rods to counter the deleterious effects of ischemia, and that RHP is facilitating this response. Moreover, the ischemic photoreceptors of *RHC mice now exhibit a significantly greater expression of several proteins regulating the ‘dark current’, including a 1.3-fold increase in the alpha subunit of the cGMP-gated cation channel protein [CNG1], 1.3-fold increases in the calcium-sensitive guanylate cyclase activating protein 1A [GUCA1A] and 1B [GUCA1B], as well as 1.4- and 1.3-fold increases in the α1 [GUCY1A1] and β1 [GUCY1B1] subunits, respectively, of soluble guanylate cyclase [GC]. None of these expression changes occurred in the ischemic photoreceptors of control mice (C1).

Comparing control ischemic retinae directly to *RHC ischemic retinae (C3) (Suppl. Fig. 5) reveals that 13 proteins in the visual transduction pathway of rod photoreceptors were expressed at levels 1.2-fold or greater between the two phenotypes, including rhodopsin kinase [GRK1] and the transducing-α–binding regulator of G-protein signaling 9 [RGS9]. Because some of these (i.e., the transducins and guanylate cyclase) are enzymes, treatment-induced changes in their abundance of more than 20% implies a very substantial modulation in visual transduction cycle function.

Collectively, these changes in the rod proteome point to the following series of biochemical responses to retinal ischemia that likely underlie the changes in their functional response to light that we measured, particularly in the rod-dominant mouse eye, with 97% of all photoreceptors being rods (*27*). With respect to the ischemic rods of F1 mice derived from control parents, the 30±4% reduction in the a-wave amplitude of the scotopic ERGs we recorded in the postischemic retina of 23 mice likely reflects ischemia-induced changes in the rod proteome that impair both the activation and the recovery phases of the visual transduction cycle. Specifically, mounting an activation (hyperpolarization) response to incoming photons is collectively impaired with less rhodopsin available to absorb them, and also less cGMP hydrolyzed per a given photonic load as a result of the lower overall catalytic power driving the closure (and resultant hyperpolarization) of cGMP-gated ion channels (secondary to ischemia-induced reductions in the levels and functionality of the co-amplifiers transducin and PDE6). The recovery phase is also impaired as a result of lower GTPase activity, secondary to ischemia-induced reductions in R9AP. However, with respect to the ischemic rods of mice derived from RHC-treated parents, the normalization of opsin and transducin-α [GNAT1] expression levels, the increased hydrolysis of cGMP secondary to increases in levels of phosphodiesterase, and the enhancement of the time of visual excitation resulting from increases in phosducin expression, are all consistent with a more stabilized activation phase of the transduction cycle. In turn, ischemia-induced impairments in the recovery phase of the transduction cycle would be countered by enhanced rhodopsin inactivation secondary to higher levels of rhodopsin kinase [GRK1], by the normalization of GTPase activity and the hydrolysis of bound GTP secondary to the greater deactivation of G-protein signaling (caused by elevations in RGS9 and S-arrestin), and by augmented expression levels of the key proteins responsible for the depolarized phenotype (dark current), most notably the catalyst guanylate cyclase. Overall, this rod proteome profile likely accounts for the significantly smaller loss of the a-wave amplitude (11±2%) in the scotopic ERGs that we recorded in the postischemic retinae of 27 *RHC mice.

## DISCUSSION

### Summary

Evidence supporting the nongenetic inheritance of disease susceptibility as a result of environmentally-induced epigenetic modifications to the parental germline continues to grow (*6–11*). Results of the present study juxtapose these findings by showing that exposure of adult mice to repetitive systemic hypoxia during adulthood – and prior to conception – protects against retinal ischemic injury in their adult F-generation progeny. Thus, not only can disease burden be inherited across generations as a result of maladaptive responses on the part of parents, but progeny can also inherit disease resilience secondary to adaptive responses on the part of their parental lineage to environments and stimuli that promote such phenotypes. Considered from an evolutionary biology standpoint, our results support the concept that short-term, single-generation enhancements in reproductive fitness occurring secondary to germline transmission of adaptive epigenetic responses can complement much longer-term, multi-generation improvements in reproductive fitness that occur as a result of Darwinian selection. Whether the adaptive response we documented is heritable by the F2 or subsequent generations will require further study. If one’s progeny are born into an environment where the potential for the same injury exists, then, from a fitness standpoint, inheriting an injury-resilient phenotype induced in parents would confer an advantage to their offspring; conversely, passing the resilient phenotype to the F2 generation and beyond would be unnecessary if the potential for these subsequent generations to experience this injury is no longer present.

To our knowledge, this is the first report of the intergenerational inheritance of an induced phenotype that protects against injury in mammals or even vertebrates. Other studies, in rodents, have demonstrated a behavioral flexibility in F1 offspring derived from F0 fathers that were subjected to maternal separation and stress during their first two weeks of postnatal life (*28*), and improvements in memory metrics in offspring from parents exposed to mentally- and physically-enriched environments (*29, 30*). However, while serving as examples of the inheritance of beneficial phenotypes, these intergenerationally modified phenotypes were not ones that protected against injury *per se*.

Whether disease or health phenotypes induced in parents secondary to environmental change are passed to their immediate or even multiple generations, at least four primary mechanisms orchestrate this entire process, each of which present fundamental biological questions. Taken ‘in order’, how systemic, environmental stress ultimately leads to germ cell reprogramming is the first. Second, what epigenetic modifications or ‘marks’ are responsible for germ cell DNA modification that survive what was long believed to be full ‘erasure’ of such marks upon fertilization? The manner in which these epigenetic marks in the zygote coordinate the ultimate differentiation of maladaptive or adaptive phenotypes in the adult, in the absence of any ecological and cultural influences on inheritance, and how these phenotypes are epigenetically maintained across the lifespan, represent the third sequential component of this larger mystery. Lastly, what are the tissue- and even cell-specific molecular signatures of these modified phenotypes? Gaining a working-level knowledge base for each of these mechanisms will take decades of concerted, interdisciplinary research. That said, as a first step to gaining a foothold on one of these unknowns, we sought to leverage the breadth and depth of proteomics to begin to understand how disease resilience, which we documented at the functional level by electroretinography, is manifested in the retina of F1-*RHC mice. For this particular question, such knowledge also carries significant therapeutic implications.

Predictably, the retinal proteome of these protected animals was strikingly distinct from that of their matched, untreated controls. Not only did we identify hundreds of differentially expressed, up- and down-regulated proteins by name, our extensive bioinformatics analyses revealed myriad biochemical pathways and networks in which they participate that were modified in the ischemic retina as a result of parental RHC, including diverse signaling and metabolic pathways not typically associated with neuroprotection. Such a pleotropic integration of phenotype and function is not unexpected given the magnitude of the functional protection we documented, as well as the complex nature of ischemic injury itself. We provided herein extensive lists of both the proteins and the key molecular pathways in which they participate with respect to both injury (C1) and protection (C2), including likely upstream regulators and downstream effectors. We also examined the RHC-induced, ischemia-resilient phenotype from the dual perspectives of how it differs from the nonischemic phenotype (C2), and how it differs from the ischemic phenotype of untreated controls (C3). All of which yielded a cornucopia of insights into the mechanisms by which intergenerational neuroprotection is established. As a result of space limitations applicable to any journal, herein we only offered mechanistic details on the basic components of a mere three of these signaling networks; however, our results are publicly available for further analyses of other pathways and proteins of potential scientific and/or clinical interest.

Dramatic changes in the levels of key signaling proteins responsible for the regulation and signaling functions of the actin cytoskeleton was one mechanism we identified and probed in detail. While not as well studied or appreciated as critical to cell function (and cell injury) as many other better known biochemical networks, our findings point to parental RHC exerting ubiquitous, functionally and structurally protective effects on a number of proteins integral to cytoskeletal dynamics (*31*), including those affecting cell shape and motility, intra- and inter-cellular protein and vesicular transport and signaling (*32*), organelle biogenesis and movement (*33*), dendritic plasticity and axon guidance, the formation of filopodia and lamellipodia, and endocytosis/exocytosis. We also provided proteomic detail regarding the proteins involved in the multi-pronged defense against systemic-level inflammatory and immune aspects of retinal ischemic injury, revealing ‘night-and-day’ differences in the expression of major acute phase response proteins (*26*) in the ischemic retinae of mice born to RHC-treated parents relative to retinae of mice descendant from untreated control parents.

From both a mechanistic and therapeutic standpoint, some of our most insightful findings related to the ischemia-protective effects of parental RHC on the proteins responsible for the visual transduction cycle. Given that the entire process of vision begins in rod photoreceptors, the lower levels of the critical transduction proteins opsin and transducing-α that we measured in ischemic retinae are likely responsible for part or all of the loss in the amplitude of the rod-generated electroretinogram a-wave that we recorded in untreated mice. Similarly, the 63% reduction in the loss of a-wave amplitude (and 83% reduction in the loss of the more distal, photoreceptor-dependent b-wave amplitude) that we measured in *RHC mice likely resulted from a remodeled rod cell proteome in which ischemia-induced losses in opsin and transducin-α are completely abrogated, and levels of the alpha subunit of the cGMP-gated cation channel, and S-arrestin, critical to establishing the dark current and the recovery phase of the transduction cycle, respectively, are increased. Such levels of endogenous plasticity in a mature tissue, and their functional consequences, would be considered impressive if uncovered in a tissue directly exposed to a given epigenetic stimulus, but the phenotypic changes we documented herein were inherited. What combination of magnitude, duration, and frequency of systemic hypoxia is required to induce the heritable, adaptive phenotype we report here is of keen interest to us, but our RHC stimulus is not sacrosanct. Other intermittent systemic hypoxia treatments (*18*), or remote limb conditioning (*34, 35*), or a pharmacological mimic (*36*) of either that ultimately modulates the expression of these proteins in a similar manner, may someday serve as a viable therapeutic for reducing the incidence of ischemic retinopathy. Also worthy of mention is that we only documented protection in the retina, and only against ischemia; given that our initial epigenetic stimulus was systemic, cytoprotective responses may occur throughout the body’s tissues, providing resilience to nonischemic injuries as well.

### Limitations

In our study, the inheritance of an induced, neuroprotective phenotype in F1 offspring resulted from exposing both parents to the epigenetic stimulus prior to mating; thus, it remains unclear whether only paternal or maternal treatment would also result in the manifestation of ischemic resilience, and to the same extent. We did not assess whether the retinae in mice from F2 or subsequent generations exhibited ischemic resilience. Regarding the aforementioned fundamental features of intergenerational epigenetic inheritance, we only interrogated the inherited phenotype at the proteomic level; measures of the underlying changes in DNA methylation, posttranslational histone modifications, and noncoding RNAs that collectively participate in epigenetics-driven changes in phenotype are still needed. Moreover, our proteomic analysis did not include the phosphoproteome (*37*), nor other post-translationally modified protein families known to be critically involved in signaling roles – these data will eventually be needed to gain a more complete picture of how the resilient proteome is established. While we used outbred mice, they were all males; sex-dependent differences may exist with respect to the mechanisms by which injury resilience is achieved. And, as alluded to above, the retinal proteomes we report here should be considered a “net” response of a set of heterogeneous changes in protein expression occurring in retinal neurons, Müller and other cells in the glial lineage, endothelial and smooth muscle cells, and even changes occurring in the blood. The time-dependency of the results of our analyses, and their implications, should also be underscored: Retinae were harvested 10 days after ischemia, so the proteome profiles reported here do not necessarily represent the ‘acute’ phase of a dynamic postischemic response, nor ones representative of ‘long-term’ recovery, but somewhere intermediate between the two. Regardless of applicable semantics, they were obtained coincident with our electrophysiologic assessments of functional outcome, supporting our causal inferences regarding the proteomic basis of resilience.

### Conclusions

The biologic complexities characterizing intergenerational epigenetic inheritance, particularly in mammals, are broad and deep. Here, we “started at the end”, by identifying the injury-resilient proteome of the functionally protected F1 retina, and dissecting a few specific signaling networks that are modified as a result of this differential expression profile. Elegant, non-trivial causal studies will be needed to parse the relative contributions of distinct proteins and pathways, and confirm their respective roles in establishing tissue protection. But for now, we hope our finding that epigenetics can modify heritability to promote injury/disease resilience will provide a compelling precedent for exploring other manifestations of inducible, beneficial phenotypes being transmitted from parents to offspring.

## MATERIALS AND METHODS

All procedures were approved by our institutional IACUC, and adhered to the ARRIVE guidelines and NIH Guide for the Care and Use of Laboratory Animals.

### Repetitive Hypoxic Conditioning

Outbred SWND4 mice of both sexes were obtained from Envigo (Indianapolis, IN) at 7-8 weeks of age, group-housed by sex (5 males per cage, and 5 females per cage), and maintained on a 12h/12h light/dark cycle with mouse chow provided ad libitum. After a two-week period of acclimation to our animal facility, our repetitive hypoxic conditioning (RHC) treatment was initiated. The mice remained in their home cages during the hypoxia exposures. Following removal of their lid covers, the cages were placed through an air-tight, sealable door into a large, gas-tight chamber (Biospherix LTD, Redfield, NY); up to 10 cages could fit into the chamber simultaneously. Thereafter, the ambient oxygen tension in the chamber was reduced to, and maintained at, 11% by flushing 100% nitrogen gas through the chamber (vent ports located opposite to inlet source) in a feedback-regulated manner, controlled by an oxygen sensor with LED readout. A two-point (electronic zero, 21%) calibration of the oxygen sensor was performed weekly. The oxygen tension was confirmed at regular intervals by a portable oxygen sensor (VTI Oxygen Analyzer, Vascular Technologies, Lowell, MA) placed within the chamber. The oxygen tension in the chamber fell from 21% to 11% in ∼5 minutes; thereafter the mice remained in the chamber for 1 or 2 hours (see treatment protocol below). At the end of the exposure period, the door to the chamber was opened, and the cages were sequentially moved into the ambient air of the housing room, lids replaced, and returned to the cage rack.

The RHC protocol we followed involved exposing the mice to 11% oxygen three times per week (M, W, F) for 16 consecutive weeks. During the initial 8 weeks, the duration of exposure was 1 h; the duration of exposure was increased to 2 h during the final 8 weeks to prevent potential habituation to the stimulus. Exposures were conducted between 8am and 12pm. Age- and sex-matched mice served as normoxic controls and were run in parallel, in that, when cages of experimental mice were moved to and from the chamber for RHC exposure, cages of control mice were also moved to and from an open shelf in the same housing room, and their cage lids removed to expose them to the normoxic room air for an equivalent period of time (1 or 2 hours).

At the gross examination level, we observed no adverse effects of the RHC stress on our F0 animals. During the initial couple of exposures to hypoxia, the mice tended to huddle together and remain relatively immobile for the duration of treatment, but by the third week of treatment, they exhibited normal cage behaviors during the hypoxic exposure. Increasing the duration of RHC to 2 hours starting on the 9^th^ week of treatment produced no further behavioral changes of note. While the F0 animals continued to gain weight, as expected, during the 16-week period of treatment, there were no significant changes in body weights between the normoxic control mice and the RHC-treated mice (data not shown). Fecundity was also unaffected by the 16-week RHC treatment; the F0 normoxic control breeders gave birth to as many litters as the F0 RHC-treated breeder pairs, and neither litter size nor male/female distributions varied between groups. Finally, the F1 progeny from RHC-treated F0 mice also appeared completely normal upon gross examination at all ages. At the cellular level, using separate cohorts of matched F0 RHC mice and F0 normoxic control mice, we found no quantitative differences in hippocampal CA1 pyramidal cell density between groups (*38*), suggesting that the intensity, duration, and frequency, as well as the overall duration of treatment that defined our repetitive hypoxic stimulus, were not acutely nor cumulatively injurious to the most hypoxic injury-sensitive neurons in the CNS.

### Retinal Ischemia and Functional Outcomes - Experimental Design

Mice received from the vendor, designated the F0 generation, were randomized to RHC or normoxic control groups, with equal numbers of males and females receiving the RHC treatment. Overall, two cohorts of F0 mice were studied; meaning, two cohorts of F0 mice derived from two separate shipments from the vendor were exposed to 16 weeks of RHC (or normoxia) and subsequently bred to generate two cohorts of F1 generation mice.

Breeding pairs of mice were established 4 days after the last hypoxia exposure by placing a single male with a single female in a clean cage. Experimental breeder groups, in which both male and female F0 mice received RHC, and control breeder groups, in which both male and female F0 mice were normoxic controls, were established. Their respective F1 progeny we termed ‘F1-control mice’ and ‘F1-*RHC mice’ (the asterisk denoting that these F1 mice were never exposed directly to our RHC stimulus). Fathers remained in the cage with the mothers and offspring until weaning, when the F1 mice were 21 days of age. Up to three F1 litters were included in the analysis, with the time between the last hypoxia exposure and conception of the given litter recorded. F1 mice were housed by sex in groups of 1-5 mice per cage. When F1 mice reached at least 4 months of age, they were subjected to 30-min unilateral retinal ischemia (see below). Electroretinography was used to quantify the extent of ischemic injury one week following ischemia (see below).

### Retinal Ischemia

Three or four days after baseline electroretinography (see below), F1 mice were anesthetized with ketamine/xylazine (100 mg/kg ketamine/10 mg/kg xylazine) intraperitoneally, and the right eye (RE) was subjected to 30 min of unilateral (right eye), normothermic retinal ischemia; the left eye (LE) was used as an internal control. Mice were thermoregulated at a body temperature of 37±0.5 °C throughout the procedure using a temp-set YSI 402 micro temperature controller (Yellow Springs Instruments) system with a rectal probe reference. Corneas were hydrated using an ophthalmic balanced salt solution. To induce retinal ischemia, a micromanipulator-mounted, 32-ga needle (attached to a 0.9% normal saline reservoir) was inserted laterally into the anterior chamber to avoid puncturing the lens. The reservoir was then elevated (178 cm) to increase intraocular pressure above systolic blood pressure, with the resultant ischemia confirmed visually in every animal by observing retinal blanching with a dissecting microscope. After 30 min, the needle was slowly withdrawn, and bacitracin-zinc neomycin ophthalmic antibiotics were applied to both eyes. Mice were monitored in a warm holding cage during their acute recovery following the procedure, and then returned to our vivarium for one week, at which point we performed postischemic electroretinography (ERG) to functionally assess outcome.

### Electroretinography

To account for small, but potentially significant diurnal/hormonal/biological variances in retinal functional activity, pre- and post-ischemic ERGs were recorded at approximately the same time of day for any given animal. Mice were dark adapted overnight prior to performing ERGs. All procedures conducted in the ERG laboratory were performed in complete darkness by the same technician using only a dim red headlamp (Princeton Tec FRED headlight, 625nm±5nm). Mice were anesthetized using intraperitoneal ketamine/xyalzine (100 mg/kg ketamine/10 mg/kg xylazine) and placed on a thermo-regulating table that maintained body temperature at 37±0.5 °C throughout the ERG recording. Scotopic response was measured using an 8-step protocol that exposed mice to a series of flashes with intensities ranging from 0.00025 cd·s/m² White-6500K (step one) to 5000 cd·s/m² Xenon (step 8). Peak amplitudes were those recorded at step 7 (250 cd·s/m² Xenon), with step 8 (5000 cd·s/m² Xenon) used only to confirm that peak response amplitudes were achieved at step 7 (step 8 not shown in figures). Three or four days after baseline ERG, mice were subjected to a unilateral retinal ischemia described in detail above; thereafter, mice were returned to our vivarium, and on postischemic day 7, scotopic ERGs were acquired again following the same 8-step, increasing flash intensity protocol as just described for baseline recordings.

### ERG Data Analysis

#### Inclusion and Exclusion Criteria

ERG waveforms were analyzed visually and in raw quantitative format exported into Microsoft Excel. Exclusion criteria included the following: Greater than 25% deviation between the RE and left eye LE in a-wave or b-wave amplitude at baseline, surgical complications, or animals becoming ‘light’ under anesthesia and requiring additional anesthesia. Any mouse that, for any reason, did not survive through the post-ischemic ERG timepoint was not included in any dataset.

ERG waveform data from the ischemic eye were normalized to those in the contralateral eye of each individual mouse as follows: The RE to LE ratios were determined for baseline and post-ischemic measurements at all steps of the protocol, and the post-ischemic RE/LE ratio was then divided by the baseline RE/LE ratio to determine, as a percentage, the overall loss of amplitude resulting from preceding ischemia.

### Statistical Analyses

Statistical analyses were conducted using STATA SE_15_ (StataCorp. 2017. *Stata Statistical Software: Release 15*. College Station, TX: StataCorp LLC) and BioVinci v 1.1.5 r2018005 (Bioturing, Inc. 2018. Data visualization software: San Diego, CA). Unpaired t-tests followed by Mann-Whitney comparison ranking were performed in GraphPad Prism (v8.3.0; San Diego, CA) to compare individual step and peak electroretinogram amplitudes with Mean+/-SEM.

### Mass Spectrometry and Proteomic Analyses

#### Sample collection and preparation

Following a single incision across the sclera with a #10-blade scalpel, the retina and lens were removed and subsequently separated from one another using sterile forceps. Retinal tissues harvested in this manner did not include Bruch’s membrane, choriocapillaris, nor eyecup/retinal pigment epithelium. Retinae were manually homogenized in 1% SDS and subsequently sonicated three times until homogeneous and sample aliquot volume was determined based on the protein concentration. 100 µg of each sample was prepared for trypsin digestion by reducing the cysteines with tris-2-carboxyethyl-phosphine (TCEP) followed by alkylation with iodoacetamide (IAA). After chloroform-methanol precipitation, each protein pellet was digested with trypsin overnight at 37 °C. Proteolytic peptides were labeled using a tandem-mass-tag (TMT)- 10plex reagent set (Thermo Scientific) according to the manufacturer’s protocol and stored at −80 °C until further use. An equal amount of each TMT-labelled sample was pooled together in a single tube and unreacted TMTs were removed using a SepPak (Waters, Ireland) under acidic reverse-phase conditions. After drying to completion, an off-line fractionation step was employed to reduce the complexity of the sample. The sample was brought up in 115 µl of 20 mM ammonium hydroxide, pH=10. This mixture was subjected to basic pH reverse-phase chromatography (Dionex U3000, Thermo Fisher). Briefly, UV-monitored at 215 nm for an injection of 100 µl and flow rate of 0.1 ml/min with a gradient developed from 10 mM ammonium hydroxide, pH=10 to 100% acetonitrile (ACN) (pH=10) over 90 min (*37*).

#### Validation using Parallel Reaction Monitoring

A Parallel Reaction Monitoring (PRM) method was developed to validate the quantification of selected proteins of interest using Skyline software (*39*); only peptides with unique sequences for the mouse proteome were considered. Figure 7 illustrates the precision of this methodology and provides validation of our overall proteomic dataset. In addition, peptides were selected for each protein-of-interest that held the following criteria: no missed trypsin cleavages, no cysteines, no methionines, and between 7-15 amino acids. Only peptides with a minimum of 6 transitions were selected for inclusion. After testing for “best flyers”, ^13^C-labelled AQUA peptides (Thermo Fisher Scientific) were synthesized to use as internal reference standards, with either a ^13^C-labelled C-terminal heavy arginine or lysine residue.

|  |  |
| --- | --- |
| Heat Shock Protein B | LFDQAFGVPR |
| Secretoglobin 2B20 | SILDYISK |
| Beta Crystallin A1 | WDAWSGSNAYHIER |

**Figure 7.**
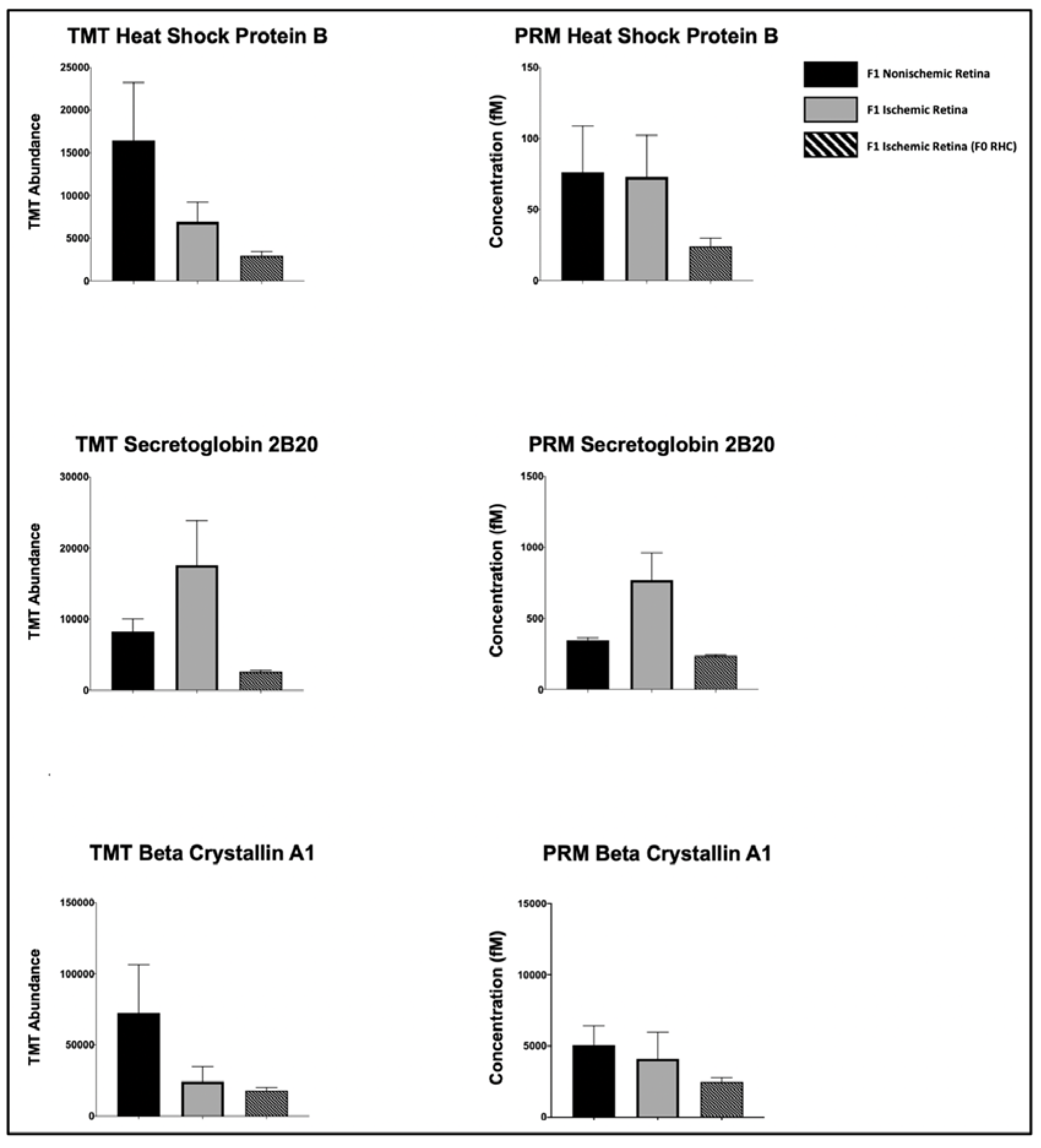
Parallel reaction monitoring (PRM) validation of a sample proteomic dataset. TMT labels provide relative quantitation across groups while PRM provides absolute quantitation. The data presented here represent the raw MS counts from samples from the following experimental groups: F1 nonischemic retina (black), F1 ischemic retina (grey), and F1 ischemic retina with F0 RHC (hatched). The TMT label abundance (left column) and respective PRM validation (right column) is displayed for the following three retinal proteins: Heat shock protein B (upper panel), Secretoglobin 2B20 (middle panel), and Beta Crystallin A1 (lower panel). Proteins were selected for validation based on robust fold-changes, or less-than-significant p-values based on initial MS runs.

Based on the protein concentration, 50µg of each sample was prepared for trypsin digestion by reducing the cysteines with DTT followed by alkylation with iodoacetamide. After chloroform-methanol precipitation, each protein pellet was digested with trypsin overnight at 37 °C.

Five µg of each tryptic digestion of all unknown mouse retina samples were “spiked” with 3.5pmol of each heavy peptide, and sample acquisition was randomized to account for any instrument drift over time. Chromatography and MS setup was as described previously. Sixty-eight-minute chromatographic runs were employed with gradient conditions as described above. A Data Independent Acquisition (DIA) was employed for the selected peptide sequences, and transitions were recorded with the following settings: Isolation m/z window of 1.4, Orbitrap resolution of 60,000, AGC target value of 5 x 105, injection time of 100ms, HCD collision of 30, and a m/z scan range of 100-1500.

Acquired PRM data were processed in Skyline. Peak integration was performed automatically and then reviewed individually for accuracy. Manual curation was based on spectra meeting the following criteria for acceptance: Consistencies in retention-time and transition retention-time, and all matching spectra having a dot product of at least 0.8 congruency with library spectra. Quantities were expressed as a ratio to known heavy peptide concentration, converted to fmols/5µg total protein, and exported into csv files for graphing.

A total of 48 fractions (200 µl each) were collected in a 96-well microplate and recombined in a checkerboard fashion to create 12 “super fractions” (original fractions 1, 13, 25, and 37 became new super fraction #1, original fractions 2, 14, 26, and 38 became new super fraction #2, etc.) (*40*). The 12 “super fractions” were then run on a Dionex U3000 nano-flow system coupled to a Thermo Fisher Fusion Orbitrap mass spectrometer. Each fraction was subjected to a 90 min chromatographic method employing a gradient from 2-25% ACN in 0.1% formic acid (FA) (ACN/FA) over the course of 65 min, a gradient to 50% ACN/FA for an additional 10 min, a step to 90% ACN/FA for 5 min, and a 10-min re-equilibration into 2% ACN/FA. Chromatography was carried out in a “trap-and-load” format using a PicoChip source (New Objective, Woburn, MA); trap column C18 PepMap 100, 5 µm, 100 A and the separation column was PicoChip REPROSIL-Pur C18-AQ, 3 µm, 120 A, 105 mm. The entire run was 0.3 µl/min flow rate. Electrospray was achieved at 2.6 kV.

#### Quantitative MS Acquisition

Isobaric TMT labels were used as reporter ions to provide unique signatures for each sample, thereby allowing for quantification of protein spectra (*41*). TMT data acquisition utilized an MS3 approach for data collection as previously described (*41*). Survey scans (MS1) were performed in the Orbitrap utilizing a resolution of 120,000. Data-dependent MS2 scans were performed in the linear ion trap using a collision-induced dissociation (CID) of 25%. Reporter ions were fragmented using high-energy collision dissociation (HCD) of 65% and detected in the Orbitrap using a resolution of 50,000. This was repeated for a total of three technical replicates. The 3 runs of 12 “super fractions” were merged and searched using the SEQUEST HT node of Proteome Discoverer 2.2 (Thermo). The Protein FASTA database was *Mus musculus*, FASTA ID= SwissProt tax ID=10090, version 2017-10-25. Static modifications included TMT reagents on lysine and N-terminus (+229.163), carbamidomethyl on cysteines (+57.021), dynamic phosphorylation of Serine, Threonine and Tyrosine (+79.966Da), and dynamic modification of oxidation of methionine (+15.9949). Parent ion tolerance was 10 ppm, fragment mass tolerance was 0.6 Da, and the maximum number of missed cleavages was set to 2. Only high scoring peptides were considered utilizing a false discovery rate (FDR) of < 1%, and only one unique high-scoring peptide was required for inclusion of a given identified protein in our results. Our mass spectrometry proteomics data has been deposited to the ProteomeXchange Consortium via the PRIDE partner repository (*42*) with the dataset identifier PXD014769.

### Bioinformatic Analyses

#### Core Analysis

Following completion of data acquisition and analysis in Proteome Discoverer 2.2, subsequent bioinformatic analyses were performed using Panther-GeneGo, STRING, and Qiagen’s Ingenuity Pathway Analysis (IPA) software. Multiple bioinformatic platforms were used to cross-reference proteins and resolve inconsistencies that may arise due to variable protein IDs across different libraries, incorrect classifications, or scheduled update differences between software packages. One initial core analysis was performed for each individual comparison (see *Results* for detail) using all high-confidence data within the dataset. Subsequent analyses were performed to refine the data using “strict” and “mild” filters, allowing for insights into highly conserved pathways as well as potential mechanistic pathways.

When annotating and applying filters for the data set uploaded to IPA, the following criteria were considered and/or applied: SEQUEST-HT scoring peptides were analyzed, and those meeting a significance threshold with a minimum abundance ratio ≥1.5 fold-change (FC) up or down, as well as statistically significant p-values were considered in the “strict” analysis. Proteins meeting a significance threshold with a minimum abundance ratio ≥ 1.2FC, and top 50% confident proteins (based on p-values) were considered in the “mild” analysis. Both the “mild” and “strict” datasets were reviewed to observe overlap of top scoring canonical pathways, upstream regulators, and biological functions. The “strict” filter was applied to identify the top 5 hits in the above-mentioned categories for each comparison; the “mild” filter was applied to procure the bioinformatics schematics shown in this manuscript. These strict/mild criteria are subjectively defined for every dataset in order to have enough protein/gene IDs to produce meaningful results, which is difficult if the filters applied are too strict, and risk too many false positives if applied filters are too mild. Each of the three comparisons was additionally analyzed in a collective “comparison analysis” in order to equate specific bioinformatic output variables across these three experimental conditions. In all bioinformatics analyses, a z-score threshold of >±2 was applied, as a p<0.05 metric for the non-randomness of directionality of a given dataset. For more detailed information on how Qiagen/IPA handles these issues in their bioinformatic analyses platform, see: http://qiagen.force.com/KnowledgeBase/KnowledgeIPAPage?id=kA41i000000L5ofCAC The bioinformatics reported herein are descriptive of the entire proteomic TMT-multiplex experiment [(2) F1-control x (2) F1-*RHC] described above, and include the core analysis summary report, and additional comparison analyses of the top Canonical Pathways, top Molecular and Cellular Functions, and top Upstream Regulators. IPA Analysis content information: Report Date: 2019-11-22, Report ID: 18557437, Content Version: 49309495 (Release Date: 2019-08-30).

## ACKNOWLEDGMENTS

Supported by NEI NIH EY018607 (JMG); NIH P30 GM106392 (JJG); Sigma Xi Grant-In-Aid-of Research (JCH); and the Louisiana Lions Eye Foundation. We also thank Nicholas Lanson Jr., Krystal Belmonte, and Moriah Harman for expert technical assistance with this study.

## AUTHORS CONTRIBUTIONS

J.M.G. conceived the study; J.C.H. and J.J.G. designed the proteomics experiments; J.C.H. and J.J.G. performed the proteomics; J.C.H., J.J.G, and J.M.G. analyzed data and interpreted experimental results; J.C.H. prepared figures; J.C.H., J.J.G, and J.M.G. drafted, edited, and revised the manuscript and approved the final version of the manuscript.

---

**Supplemental Figure 1.**
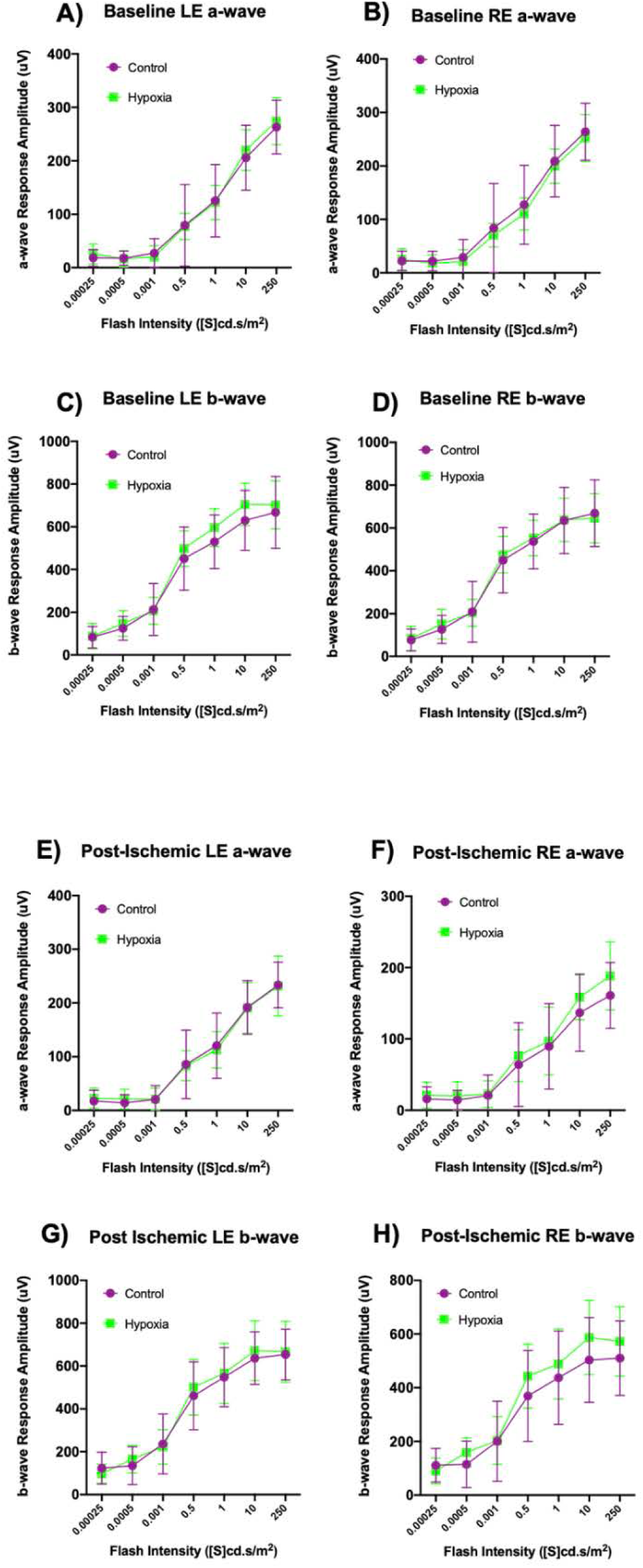
Baseline and post-ischemic scotopic electroretinography in nonischemic and ischemic eyes of F1 mice derived from control or hypoxic F0 parents. ERG a-wave **(A,B,E,F)** and b-wave **(C,D,G,H)** amplitudes are shown as a function of increasing light intensity at baseline **(A-D)** and at 10-days post-ischemia **(E-H)** for the contralateral, nonischemic left eye ([LE]; **A, C, E, G**) and the right ischemic eye ([RE]; **B, D, F, H**) for F1 mice derived from untreated F0 parents (Control, purple, [n=23]), and F1 mice derived from F0 parents treated with repetitive hypoxic conditioning (RHC) prior to mating (Hypoxia, green, [n=27]). Mean±S.D. This data is the raw data used to generate Figure 1.

**Supplemental Figure 2.**
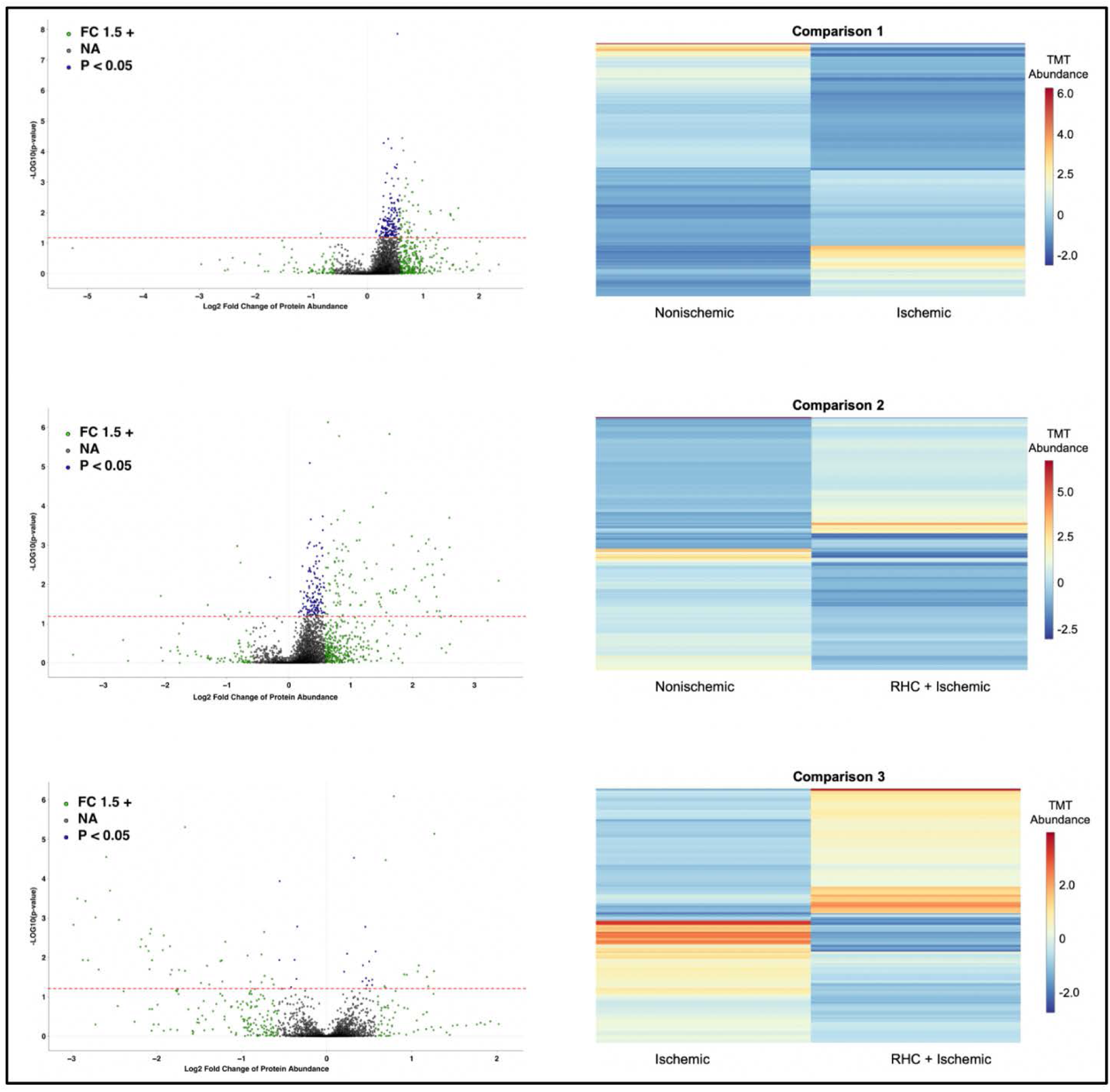
Volcano plots and heat maps reveal global differential protein expression, by experimental comparison. Comparison 1 **(upper panels)**: Ischemic retinae of F1 mice from normoxic (control) F0 parents relative to nonischemic retinae of F1 mice from normoxic (control) F0 parents; Comparison 2 **(center panels)**: Ischemic retinae of F1 mice from hypoxic (RHC) F0 parents relative to nonischemic retinae of F1 mice from normoxic (control) F0 parents; Comparison 3 **(lower panels)**: Ischemic retinae of F1 mice from hypoxic (RHC) F0 parents relative to ischemic retinae of F1 mice from normoxic (control) F0 parents. In the volcano plots (left), each point represents a unique protein, with its position on the plot based on the directional log2(fold change [FC]) of each protein (x-axis) and the -log10(p-value) (y-axis). Proteins in green were 1.5-fold or more abundant in that respective comparison. Proteins in blue are those with expression differences of p<0.05 (above dashed red line) in that respective comparison. Proteins in gray are those which did not differ significantly in either fold-change or p-value in that respective comparison. The heatmaps (right) graphically display the differential expression, for each comparison, of the top 500 up- and down-regulated proteins. The fold-change scale on the y-axis for each comparison is provided on the right y axis shows upregulated proteins in red and downregulated proteins in blue. A maximum distance with complete linkage clustering method was applied, with data centered and scaled. Each row represents the abundance of a single protein.

**Supplemental Figure 3:**
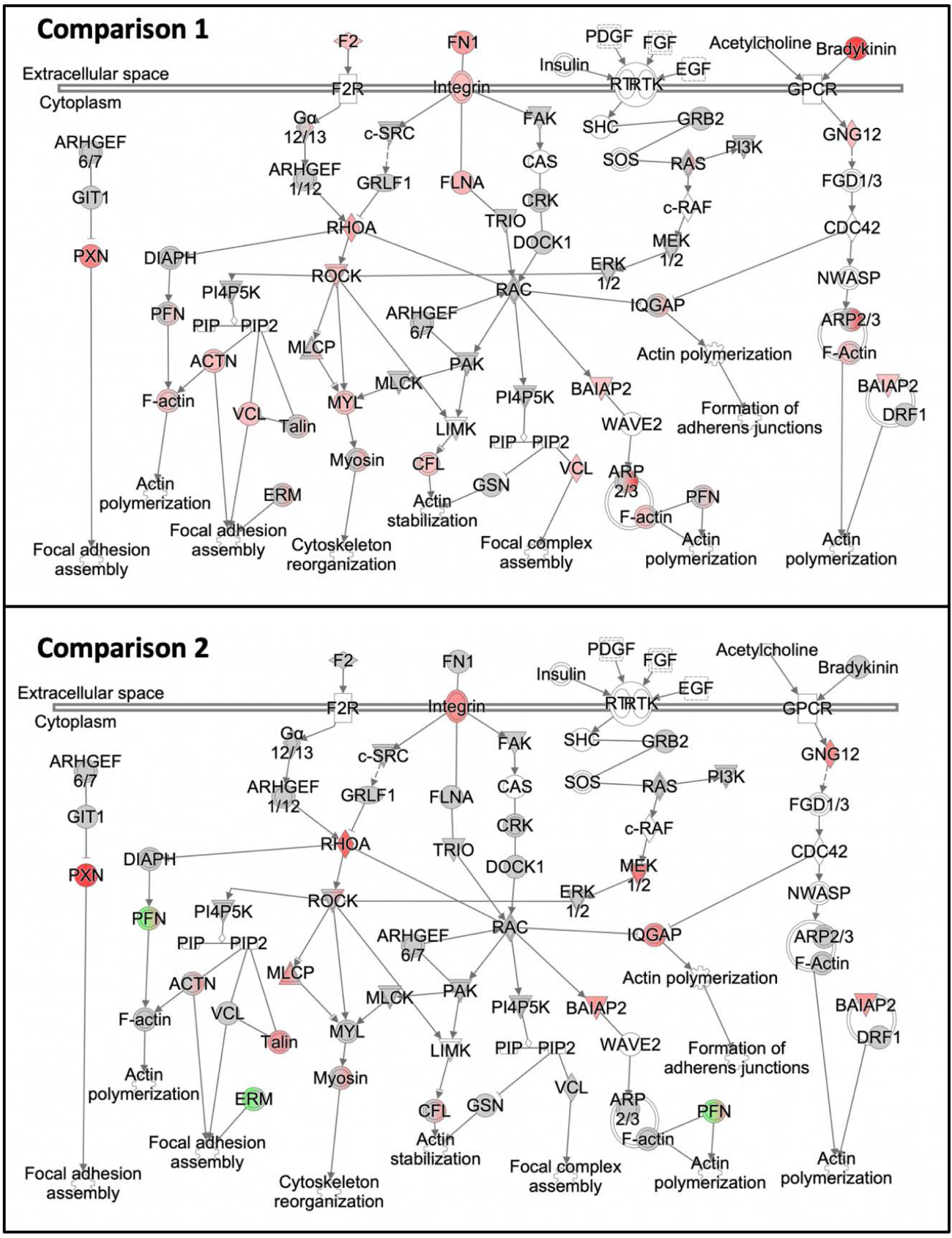
Ischemia upregulates Actin Cytoskeleton Signaling. IPA’s Actin Cytoskeleton Signaling canonical pathway is provided here for Comparison 1 **(upper panel)** and Comparison 2 **(lower panel)** (comparisons defined in main Table 1 and Supplemental Figure 3). This pathway exhibited a 6+ z-score fold-change difference between Comparison 1 and Comparison 3 (not shown). Note that Comparison 2 reveals many of the same ‘activated proteins’ (highlighted red) as Comparison 1, but to a lesser degree, and that other proteins are downregulated (highlighted green) in ischemic retinae of mice from RHC-treated parents. Proteins shaded grey exhibited some degree of differential expression in our dataset but did not meet the 1.2FC threshold applied. This figure is provided to complement Comparison 3 (Figure 3), which includes a tabulation of all of the proteins in the canonical and their respective fold changes, for each comparison.

**Supplemental Figure 4:**
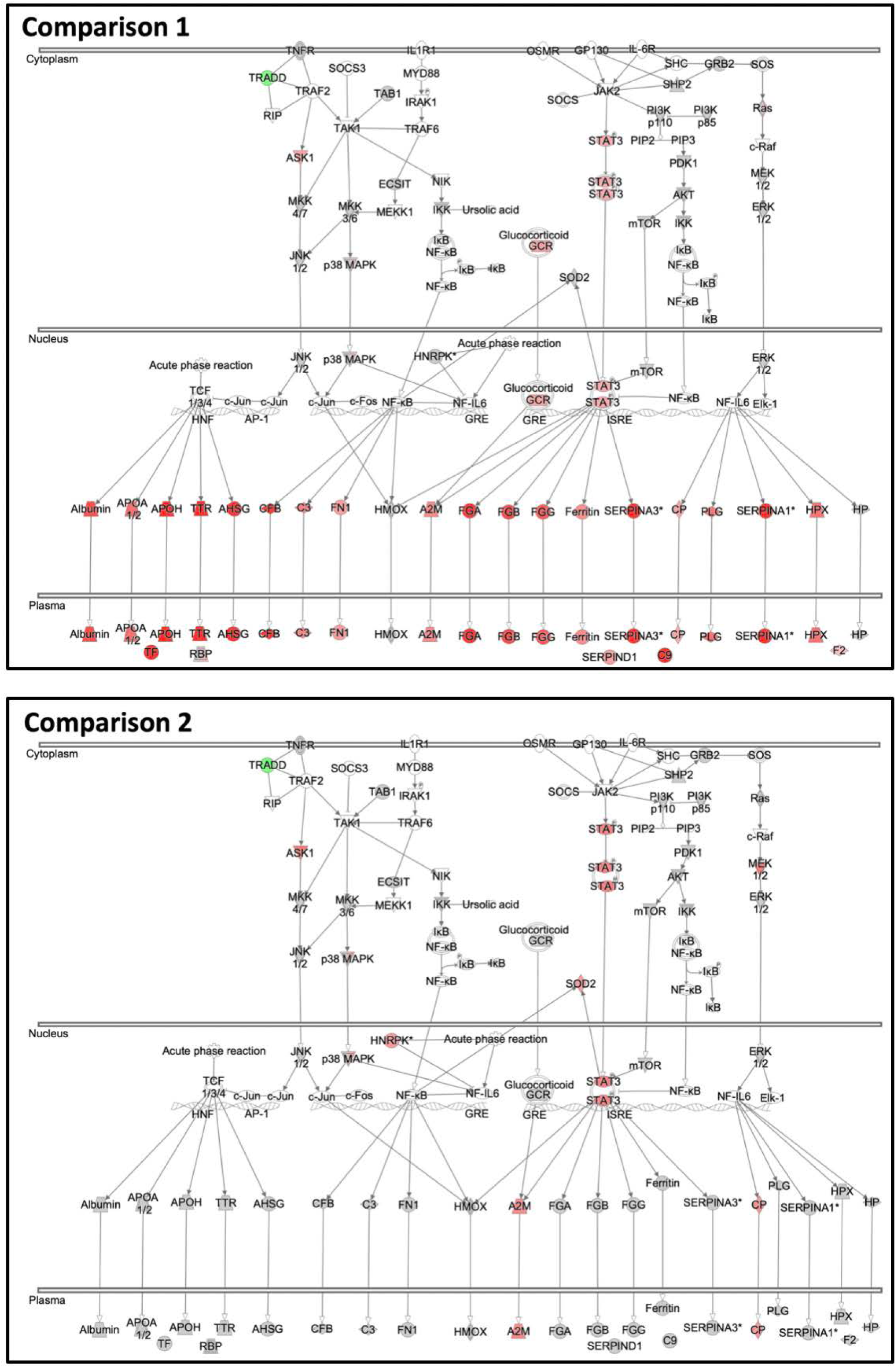
Ischemia upregulates the Acute Phase Response. IPA’s Acute Phase Response (APR) canonical pathway is provided here for Comparison 1 **(upper panel)** and Comparison 2 **(lower panel)** (comparisons defined in main Table 1 and Supplemental Figure 3). The APR was the most enriched canonical signaling pathway identified by IPA in both Comparisons 1 and 3. This pathway showed virtually no ‘activation’ in Comparison 2 at the 1.2FC threshold we applied. The APR was only observed in Comparison 3 (main Figure 3) because of the robustly opposing expression changes between the ischemic retina in mice derived from F0 parents treated with RHC and the ischemic retinae in mice from untreated F0 parents. This pathway exhibited a 4+ z-score fold-change difference between Comparison 1 and Comparison 3 (not shown). Note that Comparison 2 has many of the same ‘activated’ (upregulated) proteins (highlighted red) as comparison 1, but to a lesser degree (less red/more grey shading). Downregulated proteins are highlighted in green. Proteins shaded grey exhibited some degree of differential expression but did not meet the 1.2FC threshold applied. This figure is provided to complement Comparison 3 (Figure 4), which includes a tabulation of all of the proteins in the canonical and their respective fold changes, for each comparison.

**Supplemental Figure 5.**
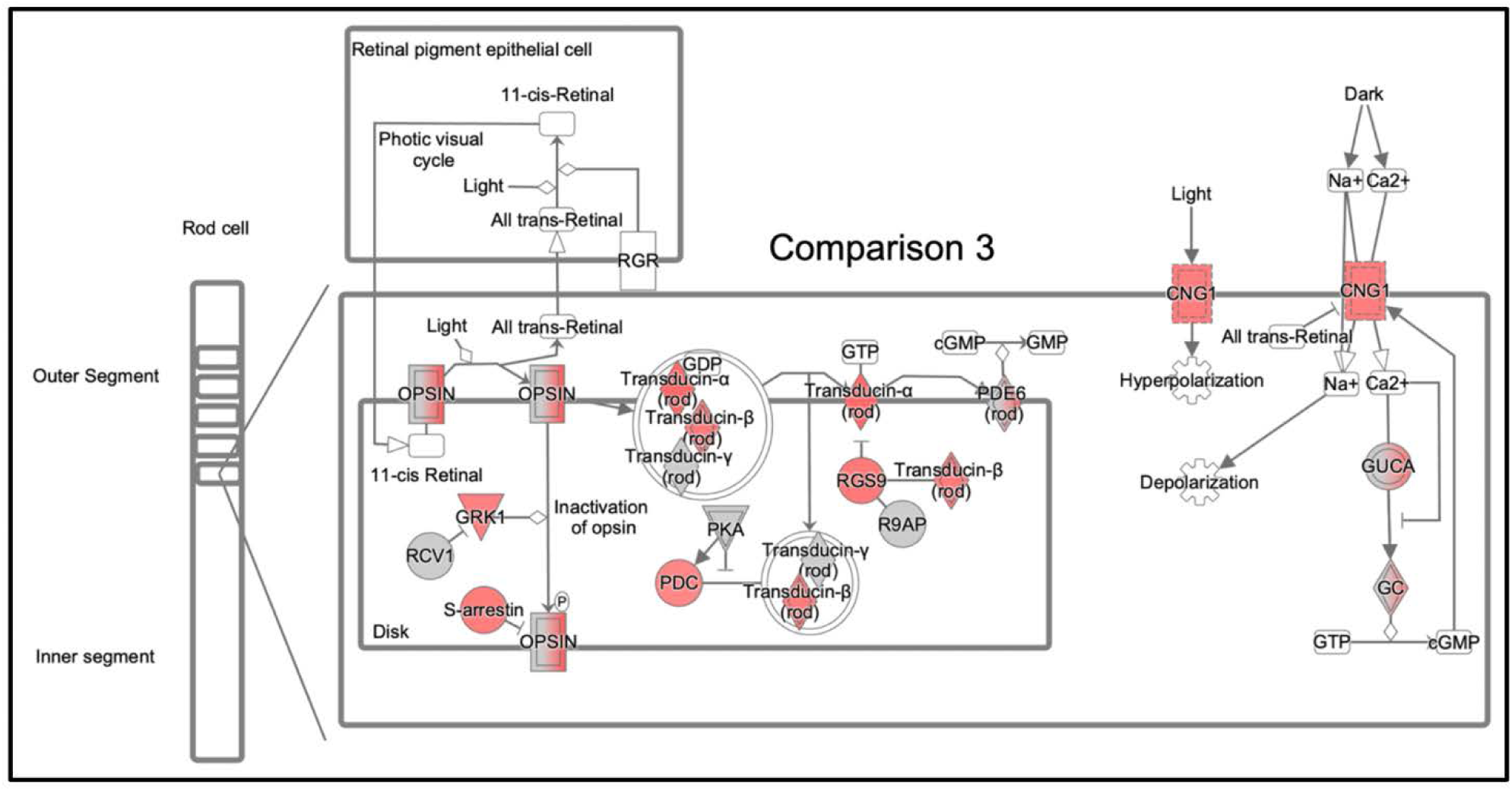
Parental RHC modulates ischemia-induced changes in rod phototransduction pathway proteins. Displayed is IPA’s version of the phototransduction canonical pathway for Comparison 3, complementing those for Comparisons 1 and 2 shown in main Figure 5. Note that many of the proteins involved in transducing light to a change in membrane potential, including some enzymes, exhibit increased expression in ischemic retinae as a result of parental RHC. A tabulation of all of the proteins in the canonical and their respective fold changes, for each comparison, is also provided in main Figure 5.

**Supplemental Table 1A.** Top 100 Differentially Expressed Proteins (abundance ratios): Comparison 1

| Protein Name | Gene ID | FC Ratio | Cellular Location | Molecular Function |
| --- | --- | --- | --- | --- |
| Actin-related protein 2/3 complex subunit 1B | Arpc1b | 4.34 | cytoplasm; cytoskeleton | protein binding; structural molecule |
| Alpha-1-antitrypsin 1-1 | Serpina1a; Serpina1d | 5.01 | ER; extracellular; Golgi | enzyme regulator activity; protein binding |
| Alpha-1-antitrypsin 1-2 | Serpina1b | 5.68 | ER; extracellular; Golgi | enzyme regulator activity; protein binding |
| Alpha-1-antitrypsin 1-4 | Serpina1d | 6.88 | ER; extracellular; Golgi | enzyme regulator activity; protein binding |
| Alpha-2-HS-glycoprotein | <u>Ahsg</u> | 5.42 | extracellular; Golgi | enzyme regulator activity |
| Antithrombin-III | Serpinc1 | 4.43 | extracellular | enzyme regulator activity; protein binding |
| Apolipoprotein A-I | Apoa1 | 3.21 | cell surface; cytosol; extracellular; nucleus | enzyme regulator activity; protein binding; transporter activity |
| Apolipoprotein A-IV | Apoa4 | 4.26 | cell surface; cytosol; extracellular | antioxidant activity; catalytic activity; enzyme regulator; metal ion binding; protein binding; transporter activity |
| Apolipoprotein C-III | Apoc3 | 3.20 | cytosol; extracellular | enzyme regulator activity; protein binding |
| Apolipoprotein E | <u>ApoE</u> | 3.28 | cytoplasm; cytosol; ER; endosome; extracellular; Golgi nucleus | antioxidant activity; catalytic activity; enzyme regulator; metal ion binding; protein binding; transporter activity |
| Arginase-1 | Arg1 | 6.15 | cytoplasm; membrane; nucleus | catalytic activity; metal ion binding |
| Beta-2-glycoprotein 1 | <u>ApoH</u> | 4.72 | cell surface; cytoplasm; ECM | enzyme regulator activity; protein binding |
| beta-crystallin A1 | Cryba1 | -1.92 | cytoplasm; nucleus | protein binding; structural molecule |
| Beta-crystallin A2 | Cryba2 | -2.08 | other | protein binding; structural molecule |
| Beta-crystallin B1 | Crybb1 | -2.86 | other | protein binding; structural molecule |
| Beta-crystallin B2 | Crybb2 | -2.50 | other | protein binding; structural molecule |
| Beta-crystallin S | <u>Crygs</u> | -2.70 | cytosol | structural molecule |
| Beta-enolase | Eno3 | -2.00 | cytoplasm; cytosol; membrane | catalytic activity; metal ion binding; protein binding |
| C2 domain-containing protein 5 | 5730419I09Rik;<br>C2cd5 | -1.79 | cytoplasm; cytosol; membrane | metal ion binding; protein binding |
| carbonic anhydrase 3 | Car3 | -1.75 | cytoplasm; cytosol | catalytic activity; metal ion binding |
| Carboxylesterase 1C | Ces1c | 5.55 | ER; organelle lumen | catalytic activity |
| Ciliary neurotrophic factor | <u>Cntf</u>; Zfp91Cntf;<br>Gm44505 | 3.48 | cytoplasm; cytosol; nucleus | protein binding |
| <u>Clusterin</u> | <u>Clu</u> | 4.65 | cell surface; cytoplasm; cytosol; ER; ECM;<br>mitochondrion; nucleus | catalytic activity; protein binding |
| Complement C1q subcomponent subunit A | C1qa | 5.82 | ECM | protein binding |
| Complement component C8 beta chain | C8b | 3.65 | extracellular | protein binding |
| complement component C9 | C9 | 5.00 | cytosol; ECM | protein binding |
| Complement factor B | <u>Cfb</u> | 6.07 | extracellular | catalytic activity |
| Cone cGMP-specific 3',5'-cyclic phosphodiesterase subunit alpha' | Pde6c | 3.57 | membrane | catalytic activity; metal ion binding; nucleotide binding; protein binding |
| Conserved oligomeric Golgi complex subunit 7 | Cog7 | -1.79 | Golgi; membrane |  |
| corticosteroid-binding globulin | Serpina6 | 3.21 | extracellular | enzyme regulator activity |
| Creatine kinase M-type | Ckm | -1.72 | cytoplasm | catalytic activity; nucleotide binding |
| DDRCK domain-containing protein 1 | Ddrgk1 | -1.82 | ER; membrane | protein binding |
| Dihydroorotate dehydrogenase (Quinone), mitochondrial | Dhodh | -1.75 | cytoplasm; cytosol; membrane; mitochondrion | catalytic activity; nucleotide binding |
| Dynactin subunit 6 | Dctn6 | 3.41 | chromosome; cytoplasm; cytoskeleton; mitochondrion | catalytic activity; protein binding |
| Exocyst complex component 6B | Exoc6b | -2.22 | other |  |
| F-actin-capping protein subunit beta | Capzb | -1.96 | cytoplasm; cytoskeleton; membrane | protein binding |
| Fascin-2 | Fscn2;<br>0610009L18Rik | -1.85 | cytoplasm; cytoskeleton | protein binding |
| Fatty acid-binding protein, epidermal | Fabp5 | -3.57 | cytoplasm | transporter activity |
| Fibrinogen alpha chain | Fga | 5.51 | cell surface; cytoplasm; ER; extracellular | metal ion binding; protein binding; structural molecule |
| Fibrinogen beta chain | Fgb | 4.15 | cell surface; cytoplasm; extracellular | protein binding; structural molecule |
| Fibrinogen gamma chain | Fgg | 3.71 | cell surface; cytoplasm; extracellular | metal ion binding; protein binding; structural molecule |
| Fibulin-1 | Fbln1 | 9.29 | extracellular | enzyme regulator activity; metal ion binding; protein binding |
| filensin | Bfsp1 | -3.86 | cytoplasm; cytoskeleton; membrane | protein binding; structural molecule |
| Gamma-crystallin A | <u>Cryga</u> | -1.89 | cytoplasm; nucleus | structural molecule |
| Gamma-crystallin B | <u>Crygb</u> | -3.13 | cytoplasm; nucleus | structural molecule |
| Gamma-crystallin C | <u>Crygc</u> | -3.45 | cytoplasm; nucleus | structural molecule |
| Gamma-crystallin D | <u>Crygd</u> | -4.35 | cytoplasm; nucleus | structural molecule |
| Gamma-crystallin E | <u>Cryge</u> | -12.50 | other | structural molecule |
| <u>Grifin</u> | <u>Grifin</u> | -3.57 | other | protein binding |
| H-2 class I histocompatibility antigen, K-B alpha chain | H2-K1 | 5.53 | cell surface; ER; Golgi | protein binding; RNA binding |
| Heme-binding protein 2 | Hebp2 | -1.79 | cytoplasm; mitochondrion |  |
| Hemopexin | <u>Hpx</u> | 3.17 | extracellular | metal ion binding; transporter activity |
| Histone H3.1 | Hist1h3i; Hist1h3a; Hist1h3g; Hist1h3h | 3.74 | chromosome; membrane; nucleus | DNA binding; protein binding |
| Ig gamma-1 chain C region, membrane-bound form | Ighg1 | 5.09 | membrane | protein binding |
| Ig heavy chain V region AC38 205.12 | IghmAC38.205.12 | 3.23 | other |  |
| Ig kappa chain C region | <u>Igkc</u> | 6.04 | membrane | metal ion binding; protein binding |
| <u>Inositol hexakisphosphate/diphosphoinositol-pentakisphosphate kinase 1</u> | Ppip5k1 | -6.25 | cytoplasm; cytosol; membrane | catalytic activity; nucleotide binding |
| <u>Interferon-induced 35 kDa protein homolog</u> | Ifi35 | 3.22 | nucleus |  |
| Isoform 2 of Eukaryotic initiation factor 4A-II | Eif4a2 | -3.33 | cytoplasm | catalytic activity; DNA binding; nucleotide binding; RNA binding |
| Isoform LMW of Kininogen-1 | Kng1 | 4.06 | extracellular | enzyme regulator activity; protein binding |
| Keratin, type II cytoskeletal 1 | Krt1 | -1.79 | cytoskeleton; membrane; nucleus | structural molecule |
| Liver carboxylesterase 1 | Ces1g | 4.30 | ER; organelle lumen | catalytic activity |
| major urinary protein 18 | Mup11; Mup18; Mup2; Mup19 | 5.24 | cytosol; extracellular; nucleus | catalytic activity; receptor activity; signal transducer; transporter |
| Methionine aminopeptidase 1 | Metap1 | 3.15 | cytoplasm | catalytic activity; metal ion binding |
| Mitochondrial 10-formyltetrahydrofolate dehydrogenase | Aldh1l2 | -1.75 | cytoplasm; mitochondrion | catalytic activity |
| Murinoglobulin-1 | Mug1 | 4.60 | extracellular | enzyme regulator activity |
| N-alpha-acetyltransferase 25, <u>NatB</u> auxiliary subunit | Naa25 | -1.75 | cytoplasm; cytosol; Golgi | catalytic activity; protein binding |
| NADH dehydrogenase [ubiquinone] 1 alpha subcomplex subunit 3 | Ndufa3 | -2.27 | membrane; mitochondrion |  |
| paralemmin-2 | Palm2 | -4.35 | membrane |  |
| Periaxin | <u>Prx</u> | -2.50 | cytoplasm; cytosol; membrane; nucleus | protein binding |
| Phakinin | Bfsp2 | -3.45 | cytoplasm; cytoskeleton; membrane | protein binding; structural molecule |
| <u>Pinin</u> | <u>Pnn</u> | -1.72 | cytoplasm; membrane; nucleus; <u>spliceosomal</u> complex | DNA binding; RNA binding |
| Plasminogen | <u>Plg</u> | 3.21 | cell surface; ECM | catalytic activity; protein binding |
| Potassium voltage-gated channel subfamily C member 2 | Kcnc2 | -2.08 | membrane | protein binding; transporter activity |
| Probable ATP-dependent RNA helicase DDX58 | Ddx58 | 3.33 | cytoplasm; cytoskeleton; cytosol; membrane | catalytic activity; DNA binding; metal ion binding; nucleotide binding; protein binding; RNA binding |
| rho guanine nucleotide exchange factor 10 | Arhgef10 | -3.45 | cytosol | protein binding |
| Rhodopsin | Rho | -2.50 | Golgi; membrane | metal ion binding; protein binding; receptor activity; signal transducer activity |
| RING1 and YY1-binding protein | Rybp | -2.08 | cytoplasm; nucleus | DNA binding; metal ion binding; protein binding |
| Serine protease HTRA1 | Htra1 | 3.65 | cytoplasm; cytosol; ECM | catalytic activity; protein binding |
| Serine protease inhibitor A3K | Serpina3k | 6.05 | extracellular | enzyme regulator activity; protein binding |
| Serine protease inhibitor A3N | Serpina3n | 10.49 | extracellular | enzyme regulator activity |
| Serine/threonine-protein kinase VRK1 | Vrk1 | -1.75 | cytoplasm; cytoskeleton; cytosol; nucleus | catalytic activity; nucleotide binding; protein binding |
| Serotransferrin | Trf | 4.79 | cell surface; endosome; ECM | metal ion binding; protein binding; transporter activity |
| Serum albumin | Alb | 3.96 | cytoplasm; ER; extracellular; Golgi; nucleus | DNA binding; metal ion binding; protein binding |
| Solute carrier family 2, facilitated glucose transporter member 1 | Slc2a1 | -1.72 | cytoplasm; cytoskeleton; cytosol; membrane; nucleus | protein binding; transporter activity |
| spliceosome-associated protein CWC15 homolog | Cwc15 | -3.85 | mitochondrion; nucleus; spliceosomal complex | RNA binding |
| syntaxin-1A | Stx1a | -2.44 | membrane | protein binding |
| T-complex protein 11 | Tcp11 | -2.13 | membrane | protein binding |
| THO complex subunit 5 homolog | Thoc5 | -4.17 | chromosome; cytoplasm; nucleus | protein binding; RNA binding |
| Transforming growth factor-beta-induced protein ig-h3 | Tgfb1 | 5.23 | extracellular | protein binding |
| Transmembrane glycoprotein NMB | Gpnmb | 3.10 | membrane | protein binding |
| Transthyretin | Ttr | 4.72 | extracellular | protein binding |
| Tubulin beta-6 chain | Tubb6 | -2.44 | cytoplasm; cytoskeleton; nucleus | catalytic activity; nucleotide binding; structural molecule |
| ubiquitin carboxyl-terminal hydrolase 8 | Usp8 | -2.17 | cytoplasm; cytosol; endosome; membrane; nucleus | catalytic activity; protein binding |
| Ubiquitin conjugation factor E4 B | Ube4b | -1.75 | cytoplasm; nucleus | catalytic activity; nucleotide binding; protein binding |
| Ubiquitin-conjugating enzyme E2 Z | Ube2z | -2.50 | cytoplasm; cytosol; nucleus | catalytic activity; nucleotide binding; protein binding |
| UPF0598 protein C8orf82 homolog | C030006K11Rik | -6.67 | mitochondrion |  |
| vitamin D-binding protein | Gc | 5.16 | cytosol; extracellular | protein binding; transporter activity |
| Vitronectin | Vtn | 4.19 | cytoplasm; ER; extracellular; organelle lumen | protein binding; receptor activity |

**Supplemental Table 1B.**
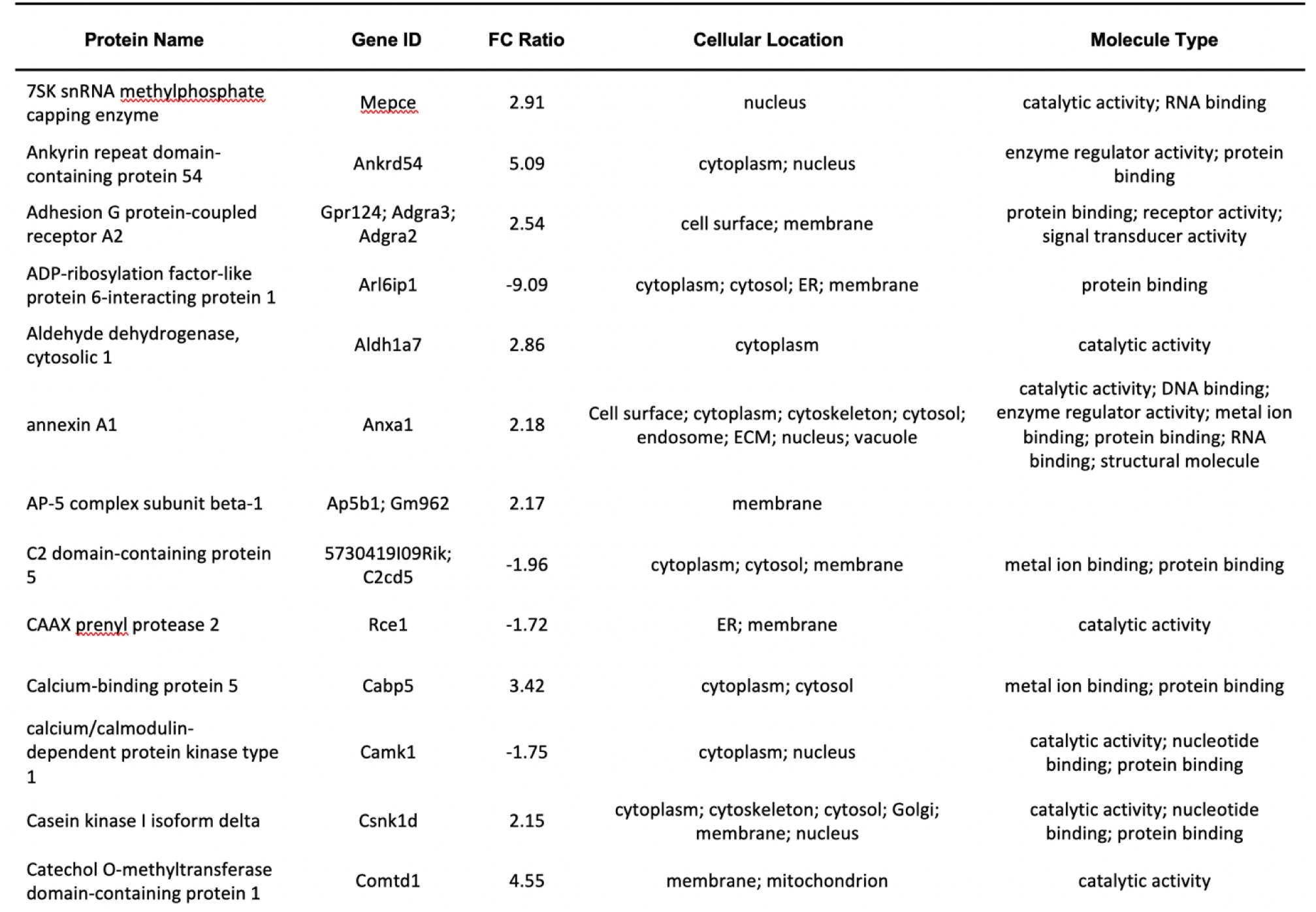
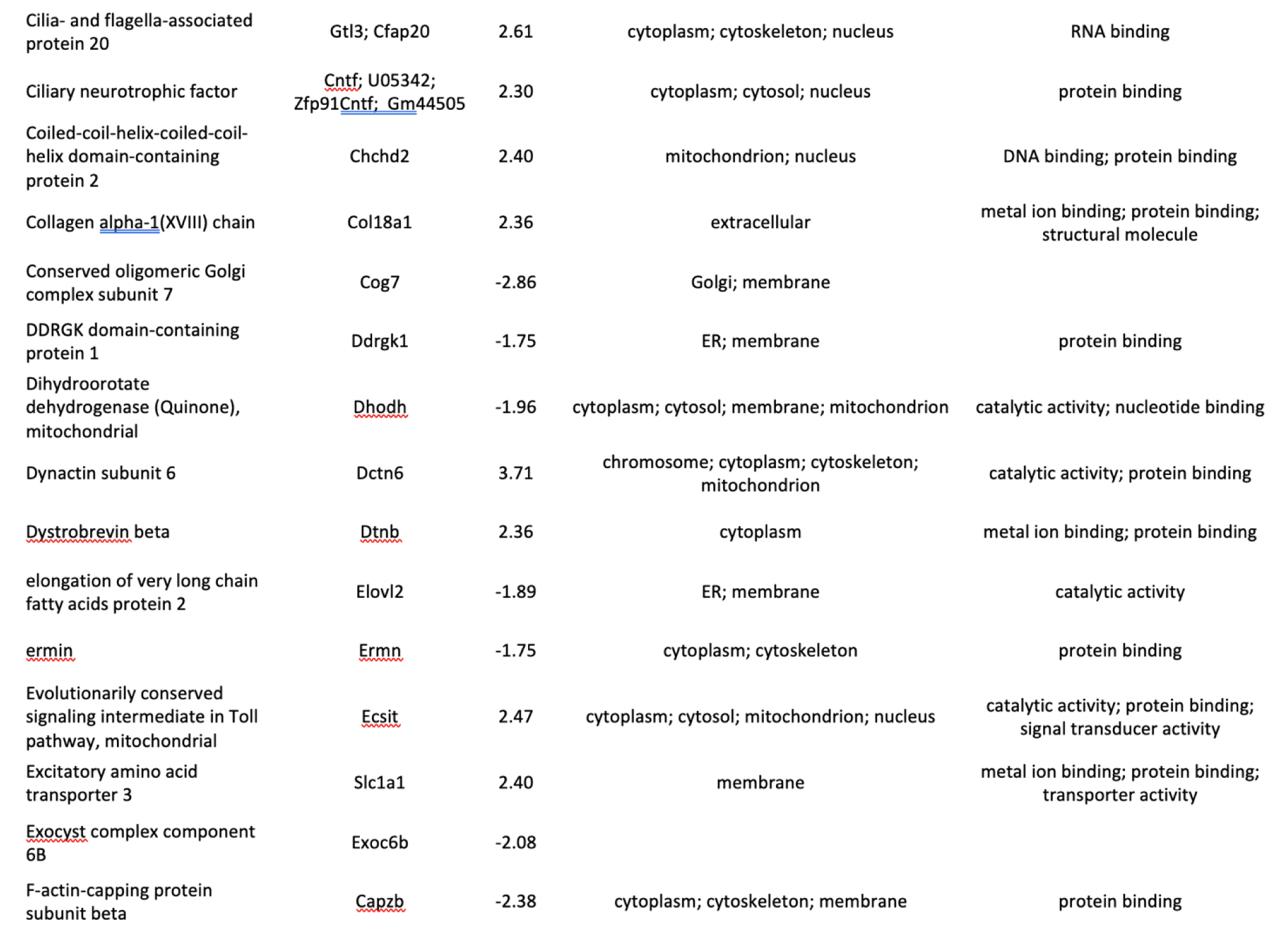
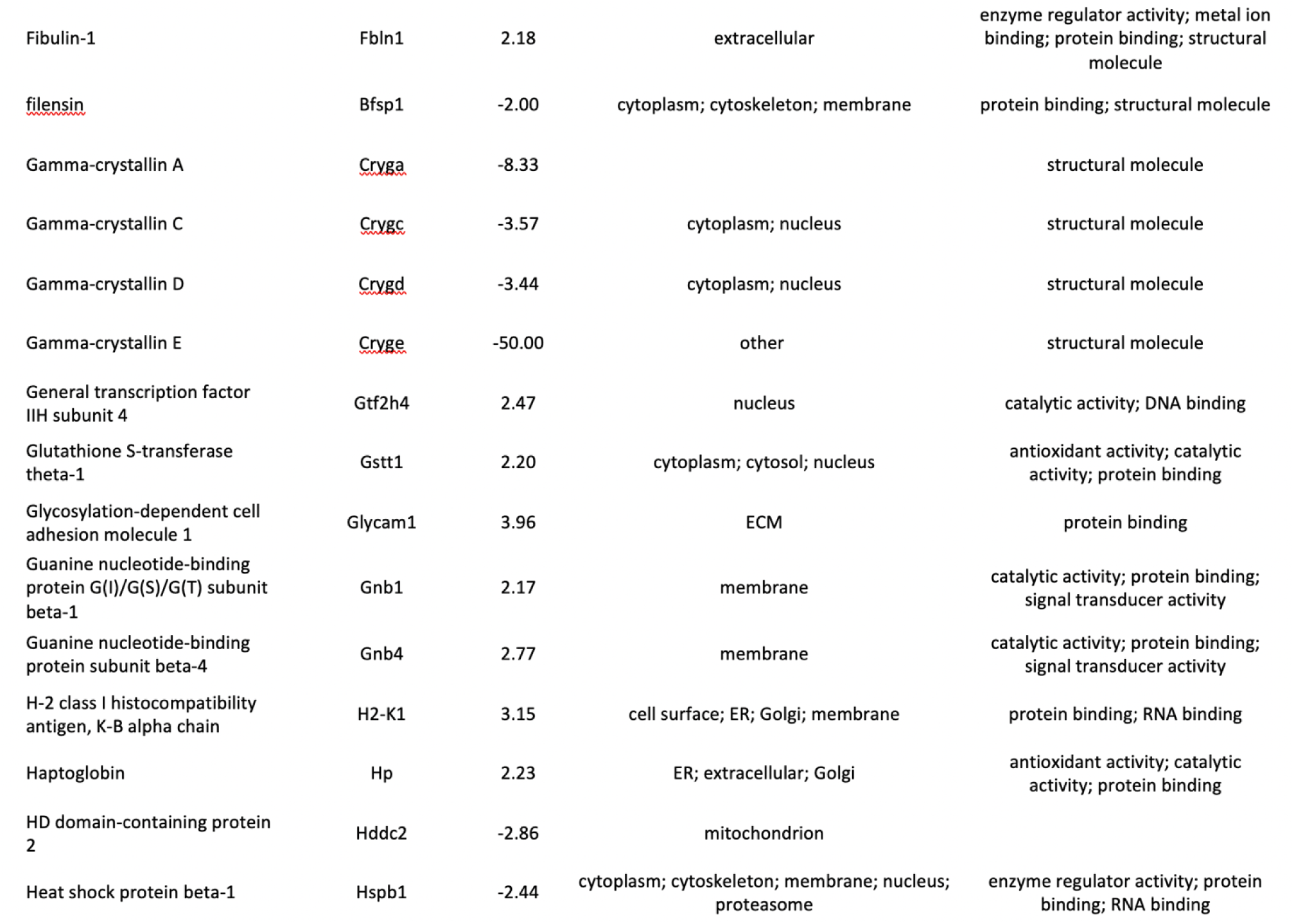
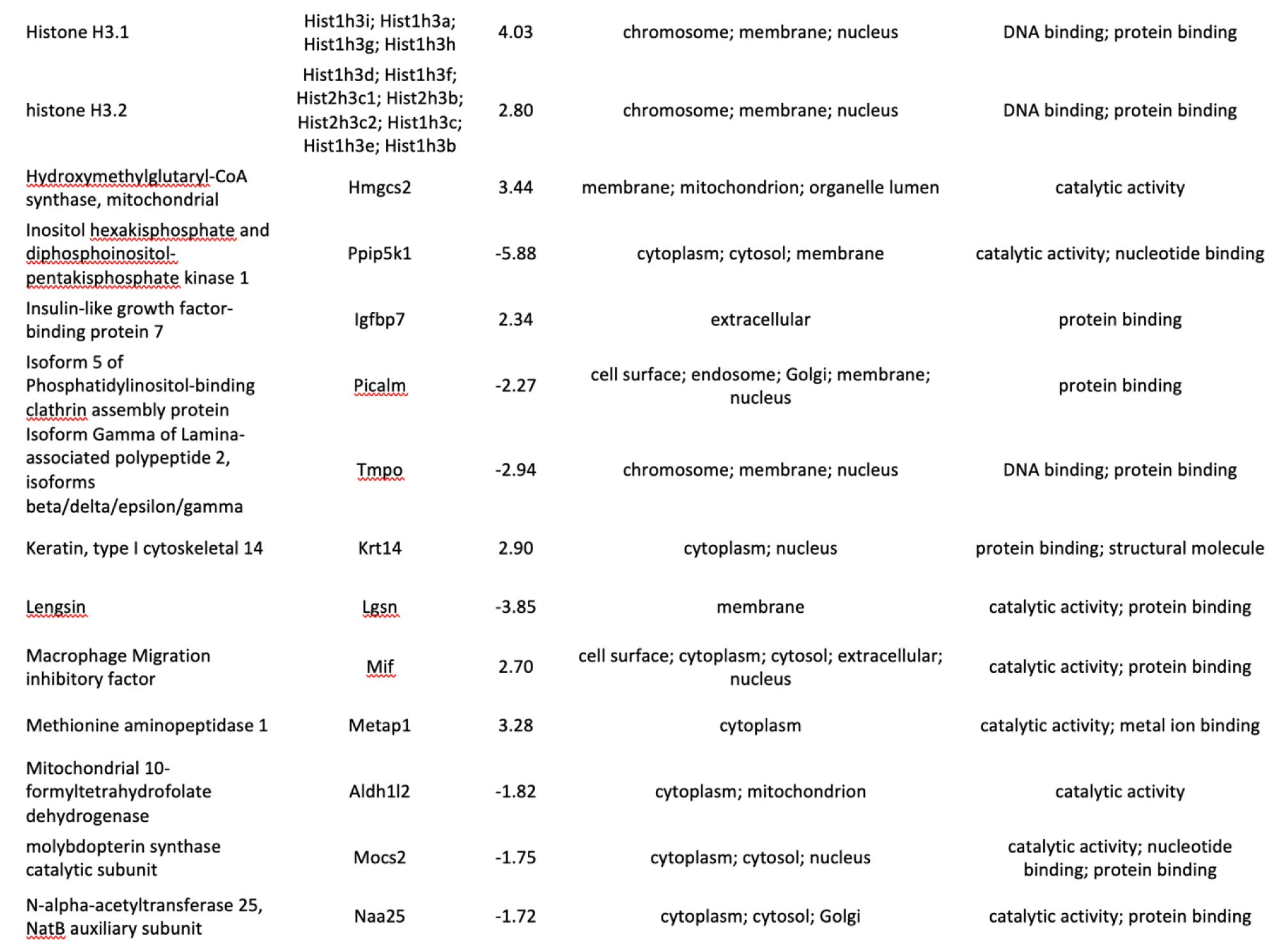
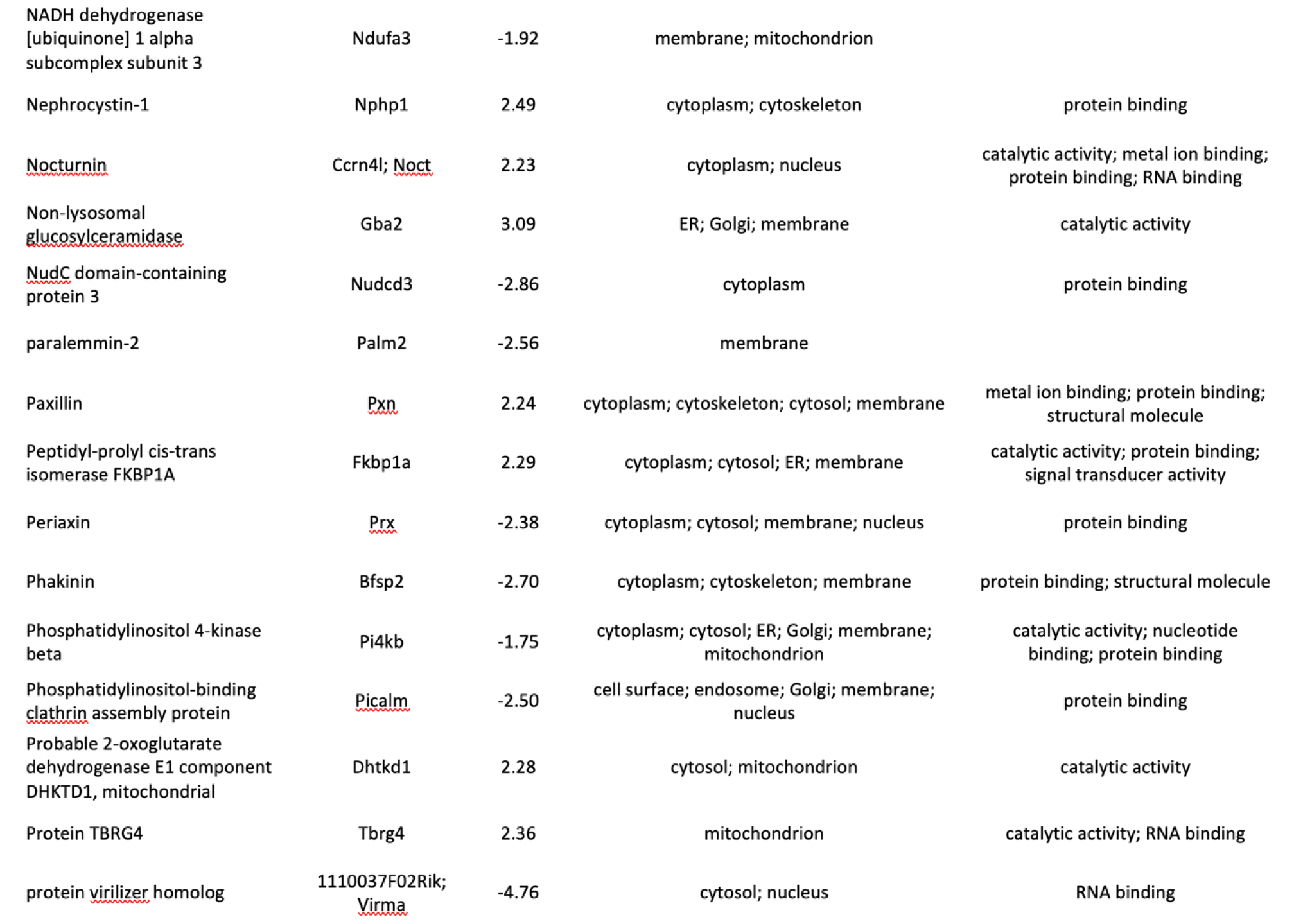
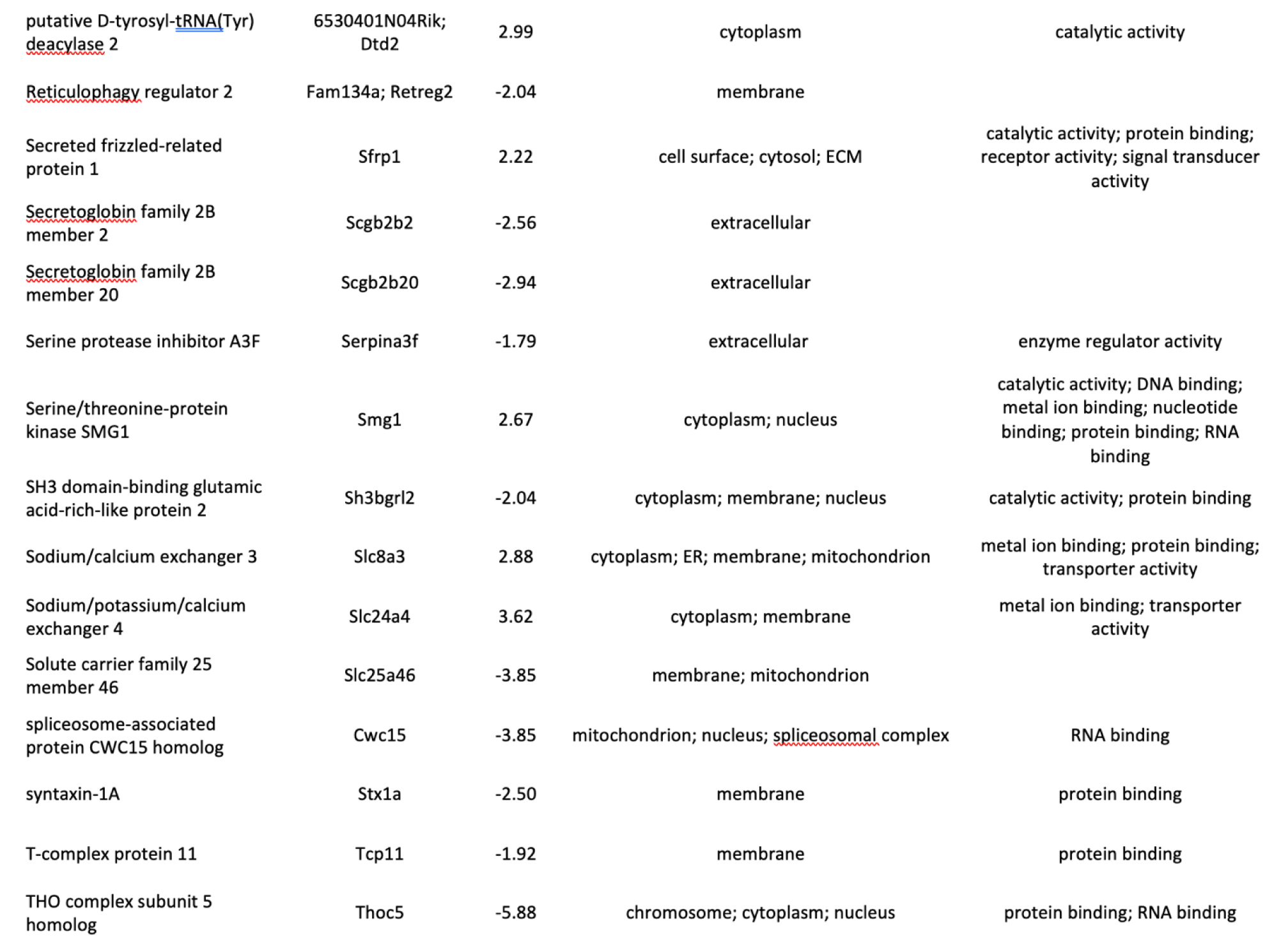
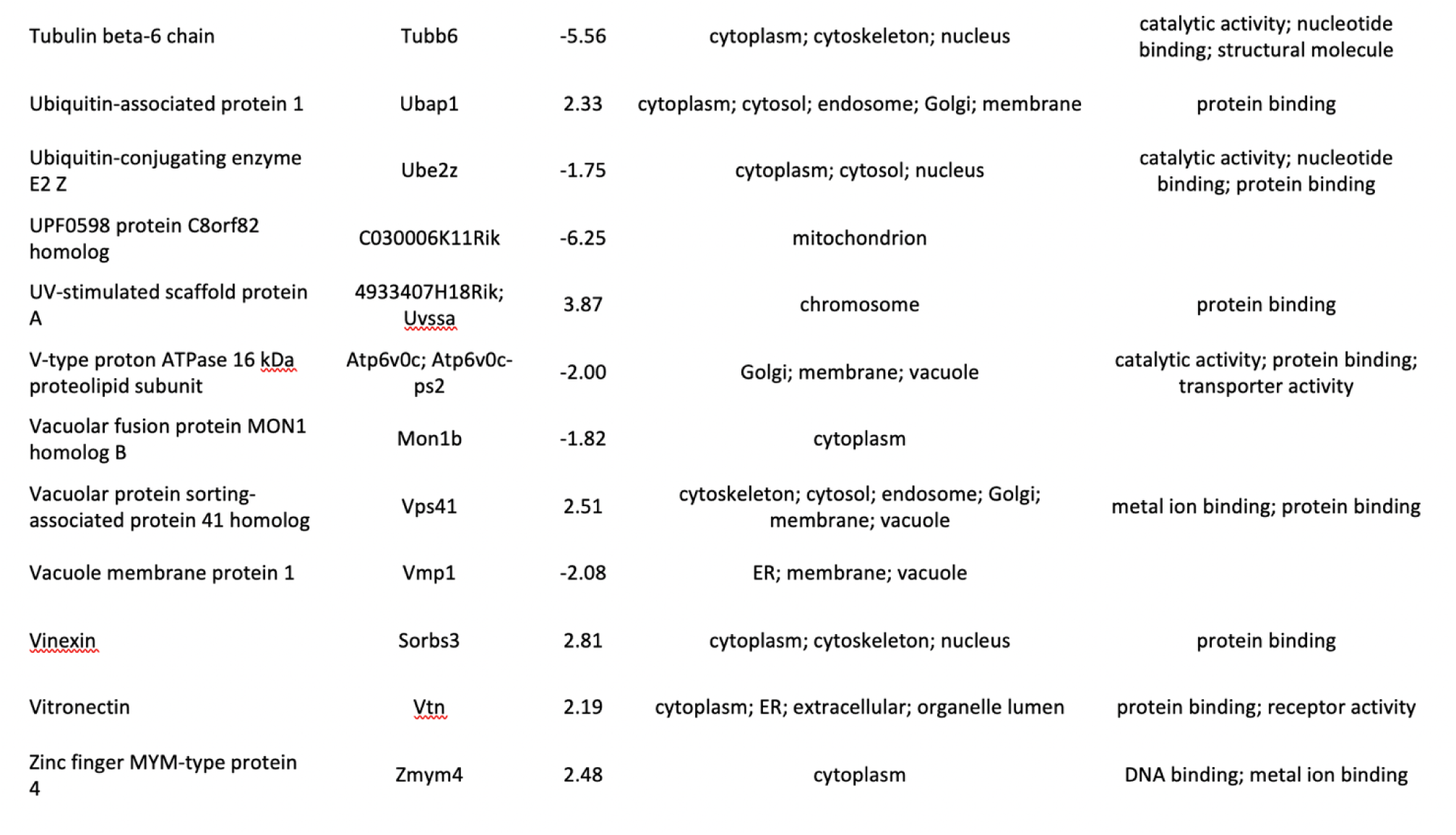
Top 100 Differentially Expressed Proteins (abundance ratios): Comparison 2

**Supplemental Table 1C.**
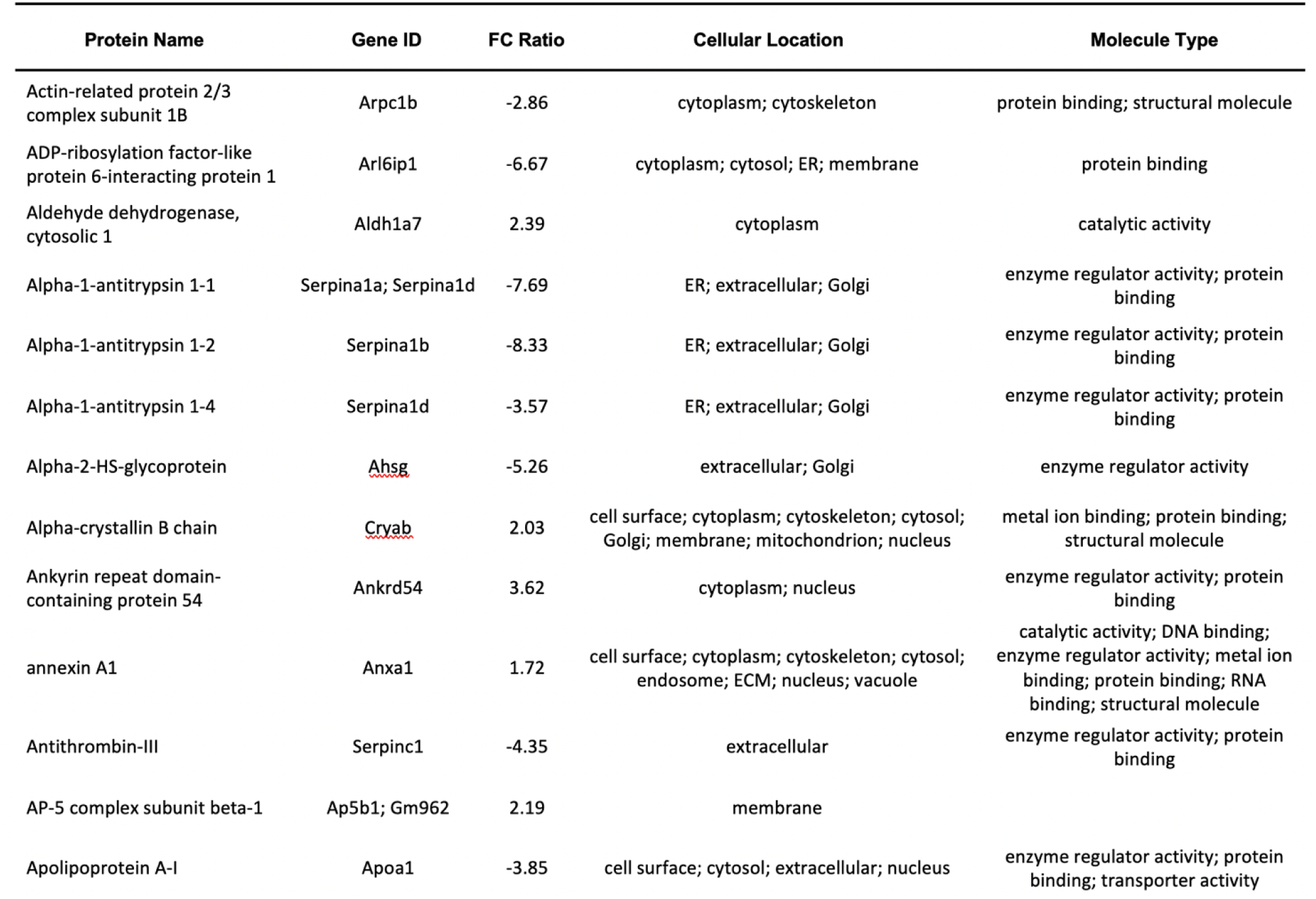
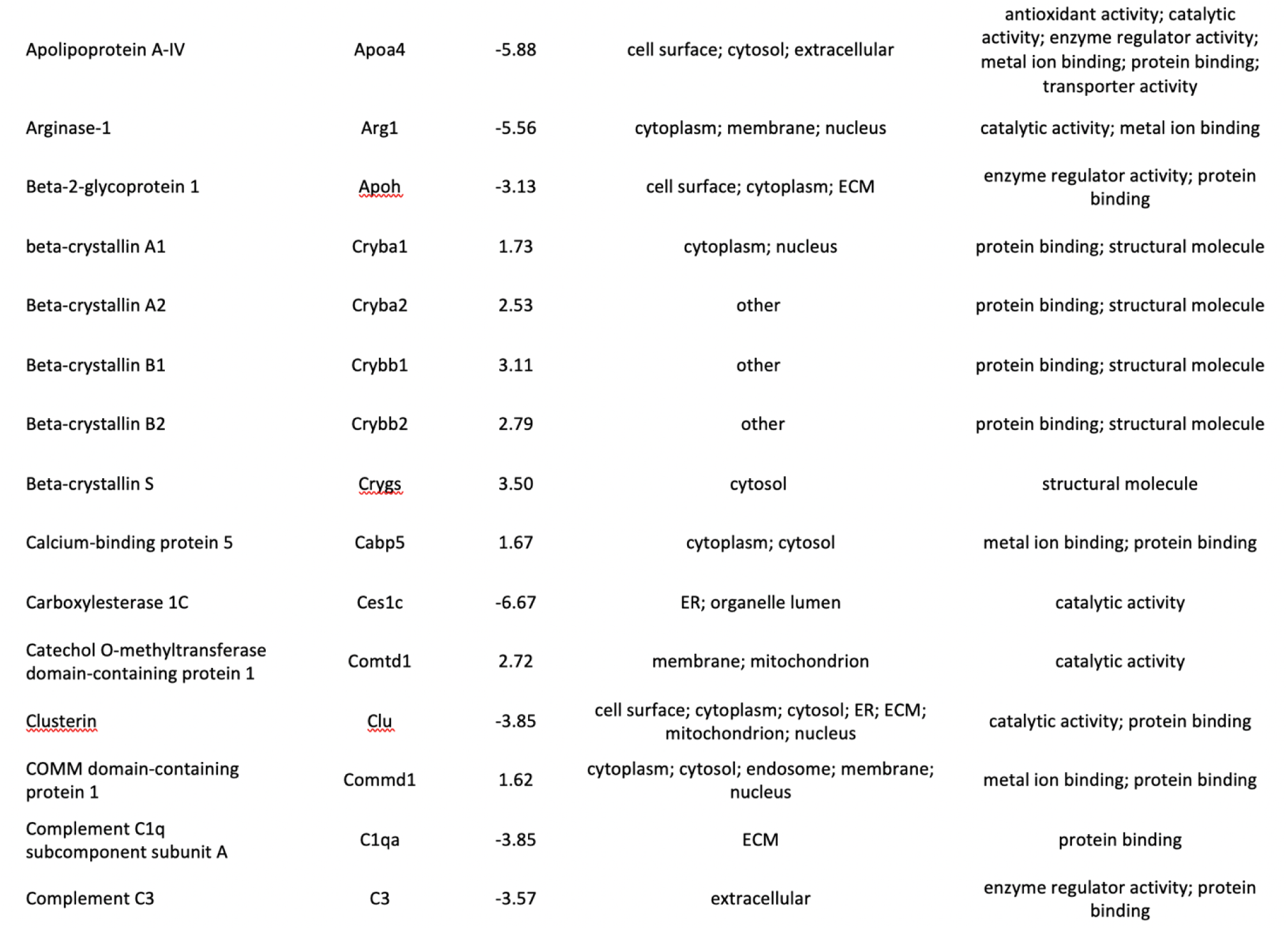
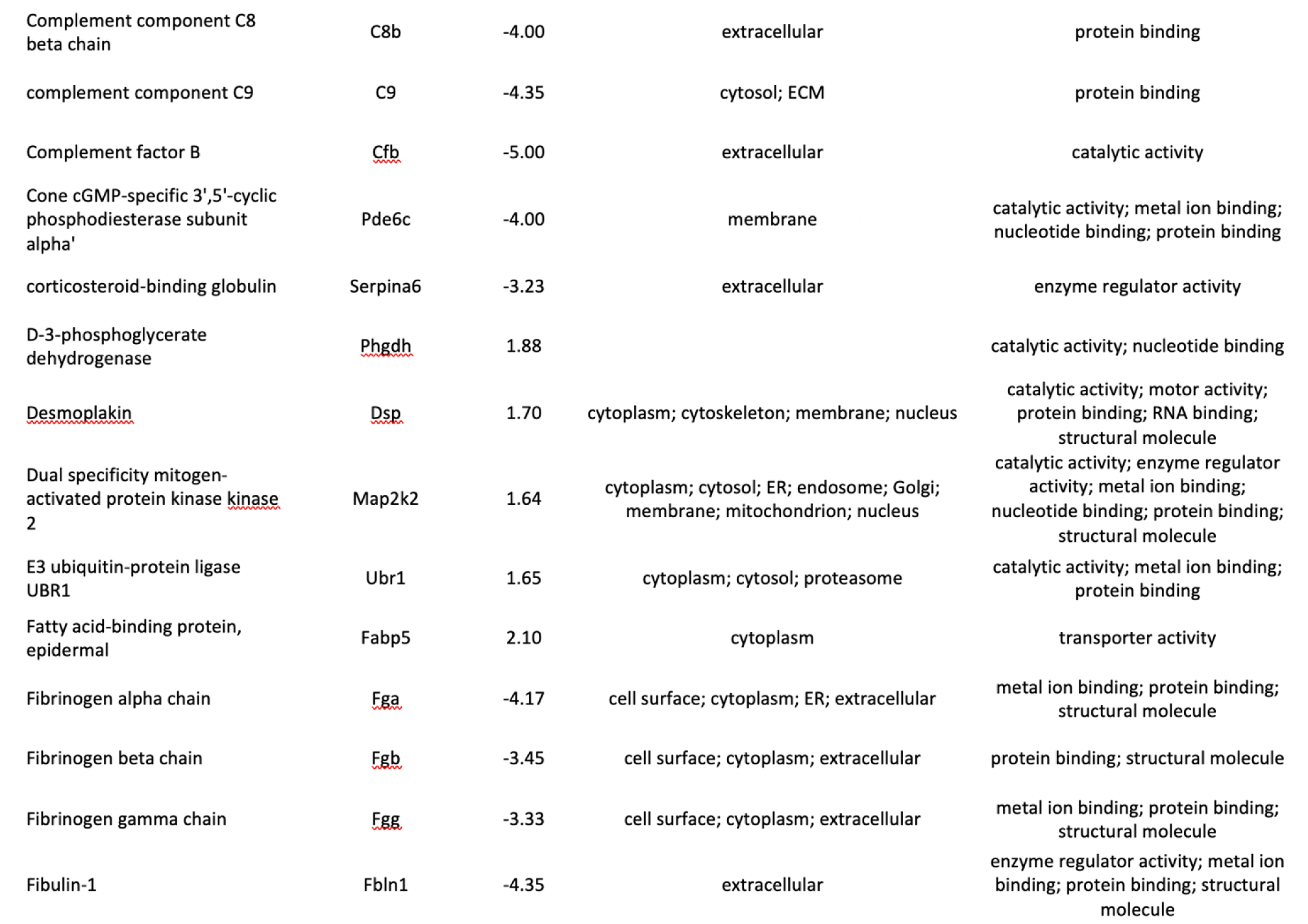
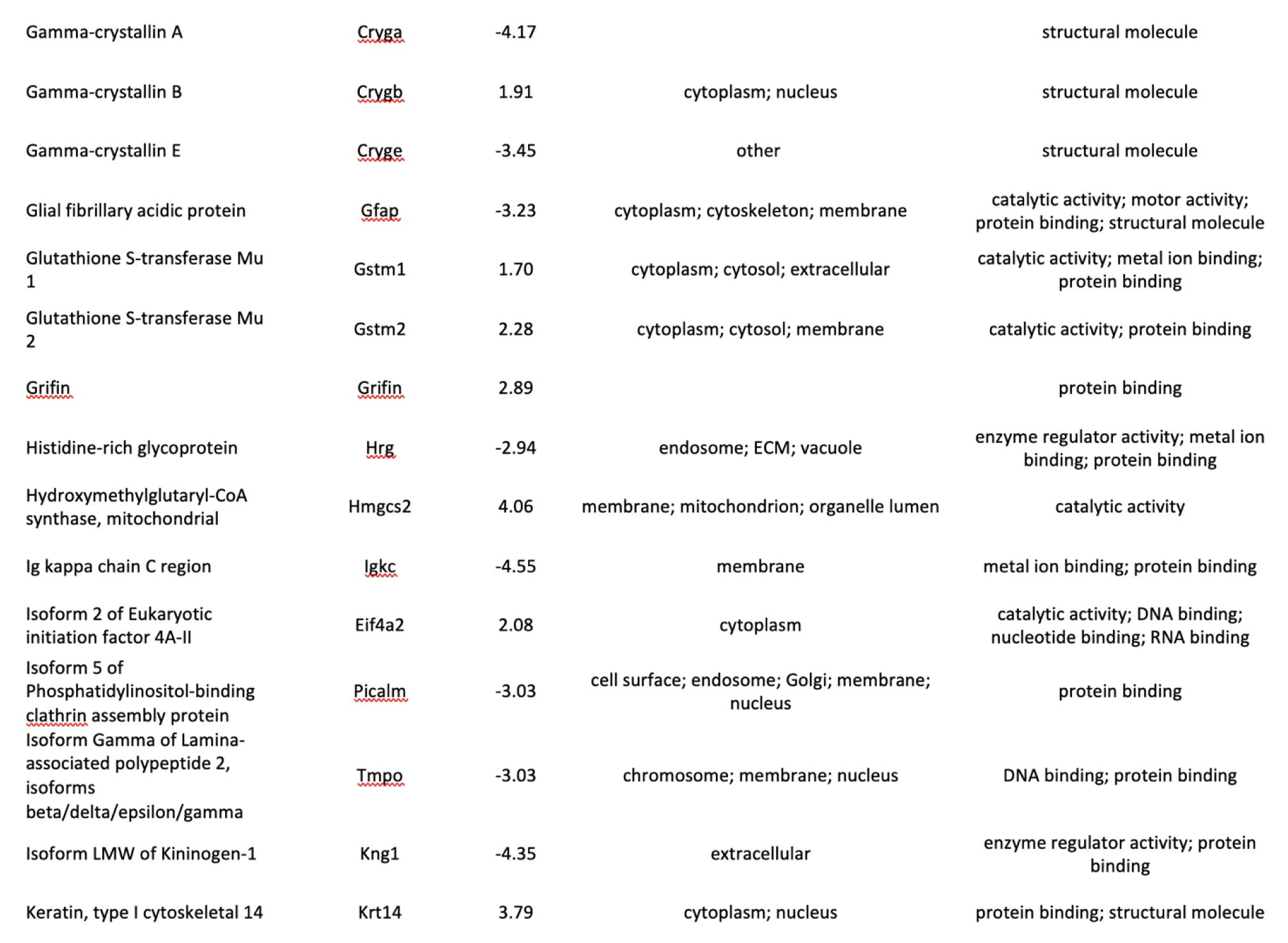
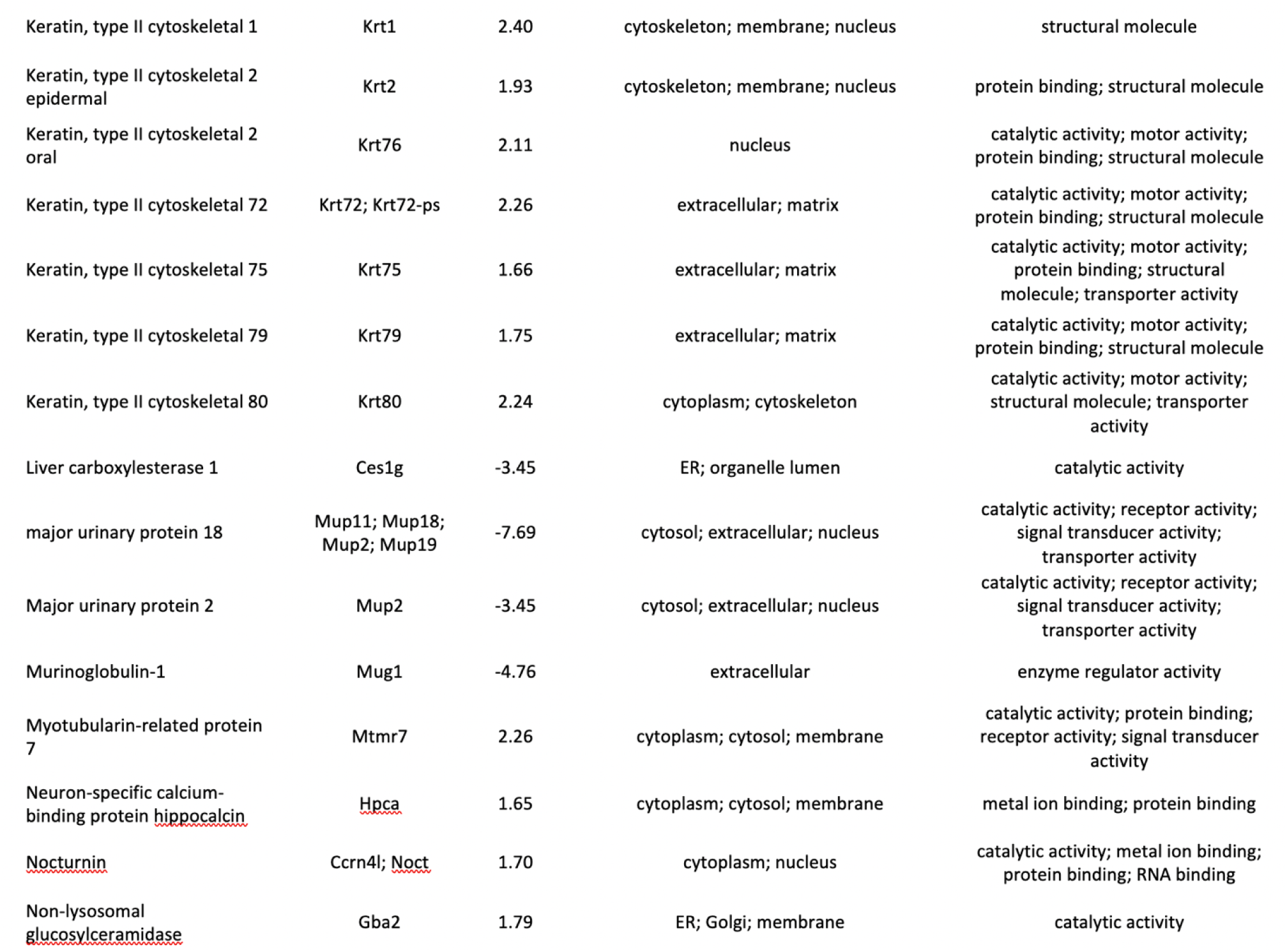
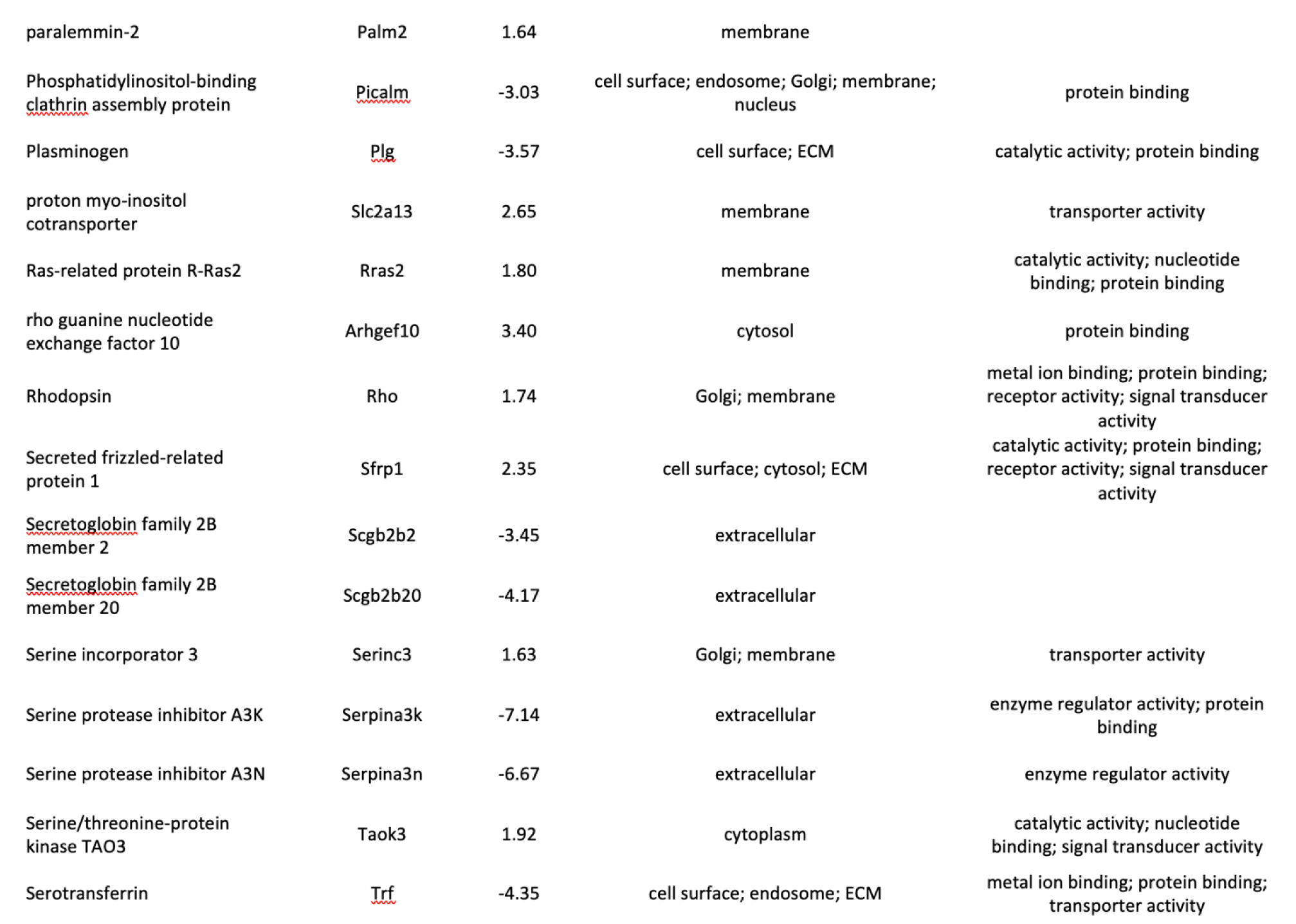
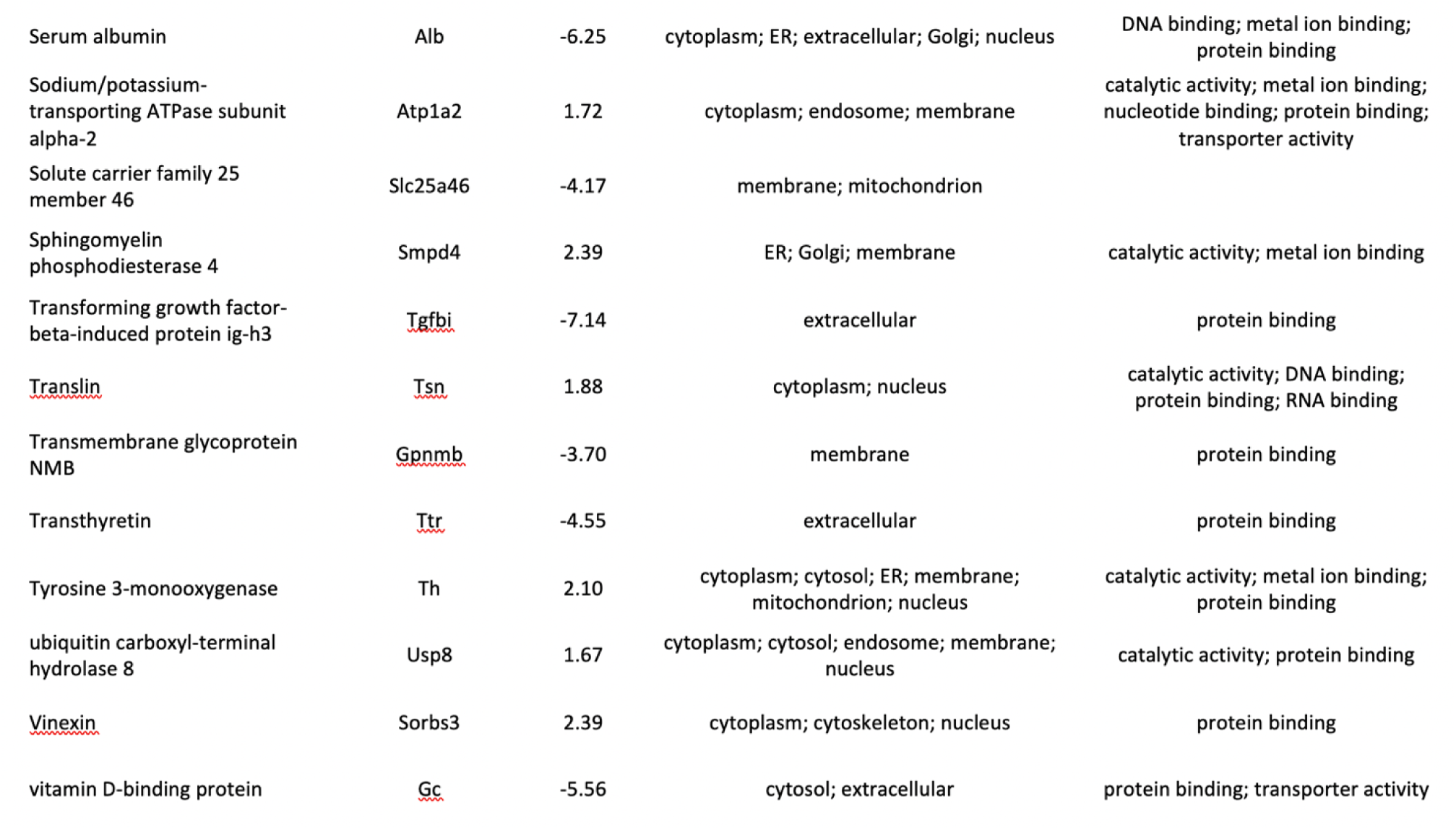
Top 100 Differentially Expressed Proteins (abundance ratios): Comparison 3

